# Enhanced human antigen-specific B cell responses using *in vitro* 3D tonsil cultures containing stromal cells

**DOI:** 10.1101/2025.09.30.679470

**Authors:** Maaike V.J. Braham, Marlon de Gast, Liubov Babii, Sabine Kruijer, Theo M. Bestebroer, Mathilde Richard, Mathieu Claireaux, S. Marieke van Ham, Jelle de Wit, Cécile A.C.M. van Els, Anja ten Brinke, Marit J. van Gils

## Abstract

The structural complexity of secondary lymphoid organs (SLOs) and their role in shaping antigen-specific B cell responses, pose significant challenges in modeling human germinal center (GC) response *in vitro*. A human 3D lymphoid model incorporating lymphoid and stromal cell types recapitulates key immune and structural features, enabling the study of antigen-specific B and T cell interactions beyond current 2D culture limitations. In this study, human tonsil cells were cultured with and without tonsil-derived fibroblastic reticular cells (FRCs) either in 2D or within a 3D PEG-4MAL hydrogel culture. Antigen-specific B cell responses in co-cultures were studied by comparing unstimulated cultures to stimulation with antigen (SARS-CoV-2 spike (S) or Influenza hemagglutinin (HA), both with or without adjuvant R848), S-nanoparticles and influenza vaccines.

Combination of FRCs with the 3D matrix significantly improved B and T cell survival and facilitated reaggregation into follicle-like structures. Antigen-specific responses were most pronounced in 3D FRC-supported co-cultures, with increasing S- or HA-specific B cell frequencies, antibody-secreting cell differentiation, and secretion of antigen-specific antibodies. Importantly, cell death and unspecific bystander activation was lowest in 3D FRC-supported cultures. Additionally, GC-associated chemokine receptors CXCR4 and CXCR5 showed distinct expression patterns on CD27⁺CD38⁺ B cells, reflecting GC-like dark and light zone organization typically observed in SLOs *in vivo*. Autologous and allogeneic FRC-supported cultures yielded comparable results, demonstrating the platform’s potential for high-throughput applications.

The 3D FRC-supported lymphoid cultures offer a physiologically relevant platform for studying human GC responses *in vitro*, supporting mechanistic research into adaptive immunity and enabling the screening of vaccine immunogens and adjuvants in a controlled setting.

## Introduction

Secondary lymphoid organs (SLOs), such as lymph nodes and tonsils, are highly complex in both structure and cellular composition. Their organization allows immune cells to interact efficiently, supported by stromal networks that guide migration and maintain tissue integrity. Upon antigen exposure, B cell follicles form specialized microanatomical structures called germinal centers (GCs), where B cells diversify their receptors through somatic hypermutation (SHM) and are selected for improved affinity (Victora and Nussenzweig 2022). This process depends on survival and differentiation signals from T follicular helper (Tfh) cells and on antigen presentation by follicular dendritic cells (FDCs) (De Boer and Perelson 2017; Wang et al. 2011). Together, these processes in the GC generate high-affinity memory B cells and plasma cells.

A defining feature of GCs is their compartmentalization into dark and light zones. In the dark zone, B cells (CXCR4^hi^CXCR5^+^) proliferate and diversify their receptors through SHM (Victora et al. 2012; Allen et al. 2004). They then migrate to the light zone, where B cells (CXCR4^lo^CXCR5^+^) encounter antigen presented by FDCs and signals from Tfh cells that drive affinity-based selection (Victora et al. 2012, 2010; Haynes et al. 2007). Cells that are positively selected may recycle to the dark zone for further diversification, or they downregulate CXCR5 and exit the GC as memory B cells and plasma cells. This cyclic shuttling between zones, coordinated by dynamic changes in CXCR4 and CXCR5 expression, ensures iterative rounds of mutation and selection that drive affinity maturation (Victora et al. 2012).

While GC organization relies on distinct zones for B cell selection, immune cell positioning within SLOs is further directed by stromal cells, including fibroblastic reticular cells (FRCs; (Lütge, Pikor, and Ludewig 2021; Acton et al. 2021; Link et al. 2007). Once considered as a mere backbone, FRCs are now recognized as active regulators of lymphocyte migration and activation (Bajénoff et al. 2006; De Martin et al. 2024). Together with FDCs, they produce chemokines that guide B and T cell trafficking (Mueller et al. 2007; Mueller and Germain 2009). Moreover, FRCs form specialized niches that support lymphocyte differentiation and help organize distinct T and B cell areas (Acton et al. 2021). By providing structural support and essential chemokines, FRCs are thus critical for the organization and function of GCs (Ahrendt et al. 2008).

Historically, animal models, particularly mouse models, have been essential in advancing our understanding of the immune system. Although mouse models have provided valuable insights, subtle differences in SLO structure, such as stromal cell composition and immune cell density, can influence GC formation and function (Mestas and Hughes 2004). Additionally, mice and humans differ in their antibody repertoires (Mestas and Hughes 2004; Schroeder and Cavacini 2010), which may influence vaccine-induced immune responses. Studying human antigen-specific immune responses, especially for vaccine development, therefore requires models that reflect key cell types and structures found in native human tissues to reliably replicate GC-like responses.

To better understand human antigen-specific B cell responses, *in vitro* models have progressed from simple 2D cultures to increasingly complex systems that incorporate multiple cell types, 3D matrices, and flow. Classical systems relied on artificial CD40L stimulation, using either recombinant human CD40L or CD40L-expressing fibroblasts in combination with varying cytokines and B cell activations (Jourdan et al. 2009; Unger et al. 2021; Marsman et al. 2022). We recently developed a synthetic human 3D *in vitro* lymphoid model to induce B cell differentiation into antibody-secreting cells (Braham et al. 2023), but it lacked natural T cell interactions and provided unrestricted CD40L stimulation in an antigen-independent manner. To study human B-T cell interactions, including selection and migration/cycling of cells, more advanced models are required. Recent studies combined 3D matrices and organ-on-a-chip technology to generate sophisticated models, in which B-T cell interactions occur in 3D structures under continuous perfusion (Goyal et al. 2022; Jeger-Madiot et al. 2024). Interestingly, none of these complex models included a stromal compartment. Additionally, the use of human peripheral blood mononuclear cells (PBMCs) often results in B and T cell populations that differ in composition and ratios from those that naturally reside in human SLOs (Bonaiti et al. 2024).

Other *in vitro* models have recently been described that use tonsil cells instead of blood-derived mononuclear cells. These models have been used to study human adaptive immune responses to varying antigen formats in the context of naturally present cell types and ratios within the tonsils (Wagar et al. 2021; Kastenschmidt et al. 2023). These tonsil models, however, are executed in more simplified 2D cultures, lacking both a 3D environment and a stromal compartment, factors described to have an essential influence on the immune responses formed in SLOs.

A 3D human lymphoid model, incorporating key tonsil-derived SLO cell types within a 3D environment, could aid the study of GC-like B and T cell responses, including their differentiation, expansion capacity, and antibody secretion and affinity. This study aims to improve current 2D *in vitro* models with the addition of a stromal compartment in a 3D environment. Tonsil-derived cells are used to assess the effect of adding FRCs, either autologous or allogeneic, as well as the introduction of a PEG-4MAL-RGD functionalized 3D matrix on T and B cell responses using different stimuli and model set-ups. Our findings demonstrate that the 3D model, with its integrated stromal compartment, enhances B and T cell survival, supports the formation of GC-like structures, and improves antigen-specific B cell responses, providing a more physiologically relevant *in vitro* model for studying the human adaptive immune mechanisms and advancing vaccine development.

## Results

### Isolation and characterization of lymphoid and fibroblastic reticular cells from tonsils at baseline

Both lymphoid (n=10) and stromal cell (n=5) fractions were isolated from tonsil donors to study the GC-like immune responses within their stromal microenvironment (Fig. 1A). The processed lymphoid fractions were directly frozen upon isolation, while the stromal fractions were first expanded. The obtained expanded stromal cell fraction was identified as fibroblastic reticular cells (FRCs; CD45^-^PDPN^+^CD31^-^ cells), expressing HLA-ABC but not HLA-DR, CD21 and CD35 (Fig. 1B).

**Figure 1.**
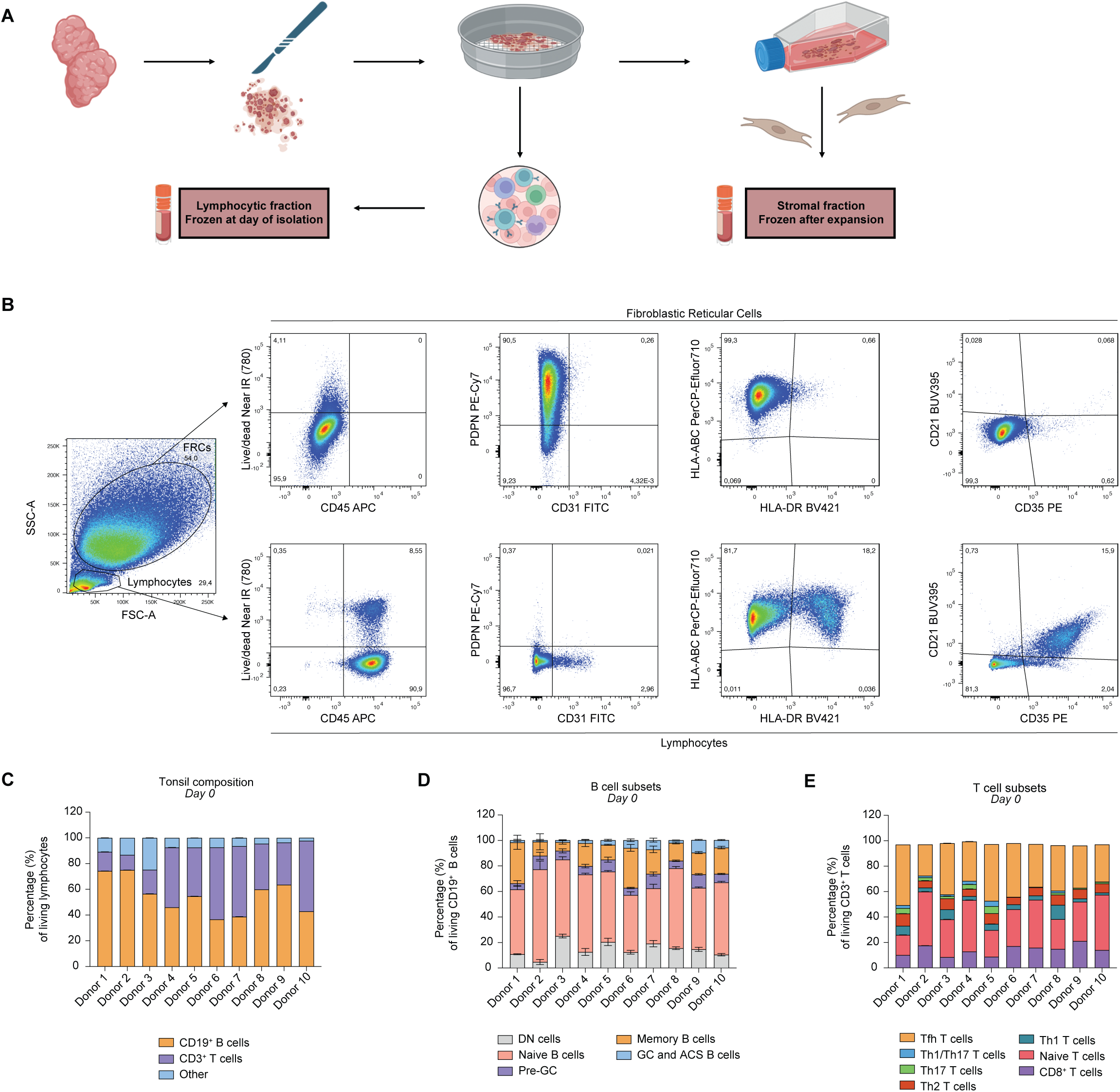
Isolation and characterization of lymphocytes and fibroblastic reticular cell fraction from tonsils. **(A)** Tonsil cells were isolated from 10 donors following tonsillectomy. Each tonsil was minced into small pieces and sieved. The isolated single-cell lymphocytic fraction was cryopreserved at day 0. The remaining tissue fragments were cultured for fibroblastic reticular cells (FRCs) isolation. **(B)** Flow cytometry plots of both the FRC and lymphocyte fractions. FRCs can be characterized as CD45^-^PDPN^+^CD31^-^ cells expressing HLA-ABC but not HLA-DR, CD21 and CD35. FRCs and lymphocytes can also be distinguished based on size and forward scatter. **(C)** Tonsil composition of each donor, expressed in percentage (%) of living lymphocytes, containing mainly CD19^+^ B cells and CD3^+^ T cells in varying ratios. **(D)** B cell subsets as a percentage (%) of living CD19^+^ B cells, containing double-negative cells (DN; CD27^-^CD38^-^IgD^-^), naive B cells (CD27^-^CD38^-^IgD^+^), memory B cells (CD27^+^CD38^-^), pre-GC (CD27^-^CD38^+^), and GC and antibody-secreting cells (GC and ASC; CD27^+^CD38^+^) in varying ratios per donor. **(E)** T cell subsets as a percentage (%) of living CD3^+^ T cells, containing CD8^+^ T cells, naive CD4^+^ T cells (CD45RA^+^), Th1 T cells (CD4^+^CD45RA^-^CXCR5^-^CXCR3^+^CCR6^-^), Th2 T cells (CD4^+^CD45RA^-^CXCR5^-^CXCR3^-^ CCR6^-^), Th17 T cells (CD4^+^CD45RA^-^CXCR5^-^CXCR3^-^CCR6^+^), Th1/17 T cells (CD4^+^CD45RA^-^ CXCR5^-^CXCR3^+^CCR6^+^) and Tfh T cells (CD4^+^CD45RA^-^CXCR5^+^) in varying ratios per donor. Data showing the mean ± SD of technical triplicates (n=2).

The lymphocytic compartment, analyzed at baseline (day 0) after freezing, consisted predominantly of CD19^+^ B cells and CD3^+^ T cells in varying ratios, with minor populations of other immune cells present (Fig. 1C, Fig. S1A). Further analysis of the B cell subsets revealed distinct populations, including naive B cells (CD27^-^CD38^-^IgD^+^), double negative (DN; CD27^-^CD38^-^ IgD^-^), pre-GC (CD27^-^CD38^+^), memory B cells (CD27^+^CD38^-^), and a CD27^+^CD38^+^ compartment containing both GC B cells (CD38^+^CD20^+^) and antibody-secreting cells (ASC; CD38^++^CD20^-^; Fig. 1D, Fig. S5.). Some donors had a mainly naive B cell compartment (up to 75% of B cells), while others had a clear memory B cell compartment (up to 37%). Similarly, T cell analysis demonstrated the presence of various T cell subsets, including Th1, Th2, Th17, and a substantial portion of the GC essential Tfh cells, which constituted up to 50% of the total T cells (Fig. 1E, Fig. S1B.).

To study B and T cell responses to SARS-CoV-2 virus and influenza H1N1 virus antigens, presence of antigen-specific B cells to both viruses were characterized at baseline, focusing on their specificity for SARS-CoV-2 Spike (S) and influenza H1N1 hemagglutinin (HA; Fig. 2A). Antigen-specific B cells were detected in all donors for both antigens, at varying frequencies and phenotypes (Fig. 2B). Antigen-specific B cells were mainly found in the class-switched double negative (CD27^-^IgD^-^) and classical memory compartment (CD27^+^IgD^-^), mainly being IgG^+^ cells (Fig. 2C-D). The observed variability in antigen-specific B cell populations underscores considerable donor-to-donor differences in immune histories and highlights the importance of analyzing baseline cells as the starting point for subsequent *in vitro* cultures.

**Figure 2.**
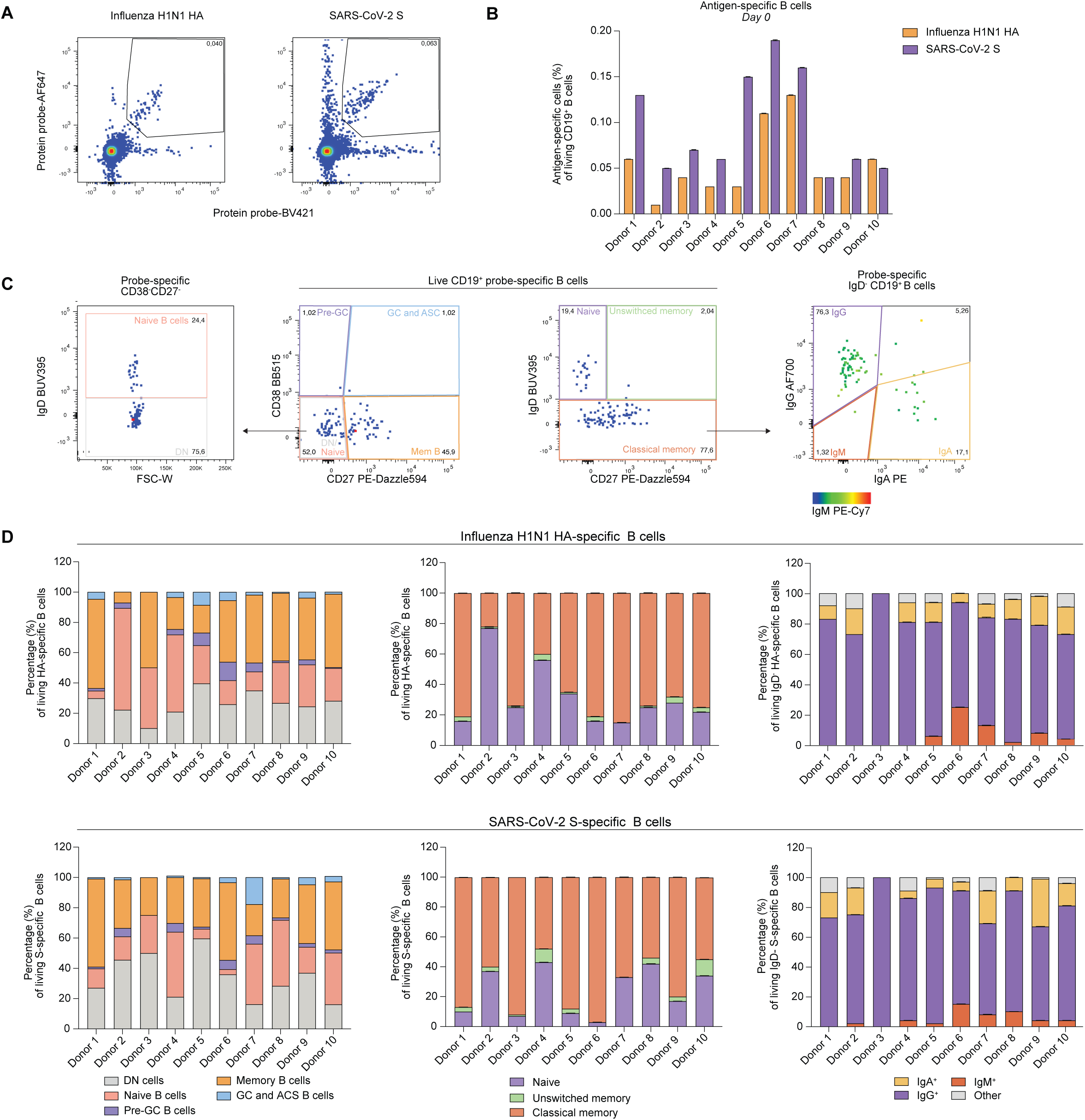
Baseline characterization of influenza H1N1 hemagglutinin (HA)-specific B cells and SARS-CoV-2 wild type (WT) spike-specific B cells. **(A)** Flow cytometry plots of protein-specific B cells gating. Viable CD19^+^ B cells were considered protein-specific when double-positive for the used protein probes in two fluorescent channels. **(B)** Influenza H1N1 HA-specific B cells and SARS-CoV-2 WT spike-specific B cells as a percentage (%) of living CD19^+^ B cells per donor. **(C)** Flow cytometry plots illustrating the gating strategy for live CD19⁺ probe-specific B cells. CD27 and CD38 expression (middle left plot) was used to identify naïve/double-negative (DN; CD27⁻CD38⁻), memory (CD27⁺CD38⁻), pre-germinal center (pre-GC; CD27⁻CD38⁺), and germinal center/antibody-secreting cells (GC/ASC; CD27⁺CD38⁺). From the CD27⁻CD38⁻ subset, DN and naïve B cells were further distinguished based on IgD expression (left plot). From the live CD19^+^ B cell population, IgD and CD27 expression (middle right plot) was used to identify naïve (IgD⁺CD27⁻), unswitched memory (IgD⁺CD27⁺), and class-switched memory (IgD⁻CD27⁺) B cells. Class-switched memory B cells were subsequently analyzed for isotype expression and subdivided into IgM⁺, IgG⁺, or IgA⁺ subsets (right plot). **(D)** Characterization of viable influenza H1N1 HA-specific B cells (top row) or viable SARS-CoV-2 WT spike-specific B cells (bottom row) in terms of CD27/CD38 expression (left), IgD/CD27 expression (middle) and IgM/IgG/IgA (right) into the populations elaborated upon (C), shown as a percentage (%) of total viable protein specific B cells. Data showing the mean.

### Setup of an *in vitro* lymphoid model, enhancing traditional 2D cultures by incorporating both FRCs and a 3D matrix resulting in cultures resembling SLO structures

To establish a culture system that closely mimics SLOs, we aimed to enhance traditional 2D cultures by incorporating FRCs and a 3D matrix. The 3D matrix used is a synthetic PEG-4MAL hydrogel (5% (w/v)), free of any animal-derived components, cross-linked with a protease degradable cross-linking peptide GCRDGPQGIWGQDRCG (GPQ-W) peptide and cell adhesive peptide GRGDSPC (RGD), which mimics the minimal integrin-binding motif found in extracellular matrix proteins (Braham et al. 2023) (Fig. 3A). The incorporated RGD peptides promote integrin-mediated adhesion, allowing cells to attach and spread effectively within the 3D environment, thereby enhancing cellular interactions in a more physiologically relevant manner. Within these hydrogels, the tonsillar lymphocytic and FRC fractions were mixed, forming cultures that were visualized using fluorescent confocal microscopy. Imaging revealed reaggregation and close interactions between B and T cells (Fig. 3B). The FRCs form structural niches/follicles within the hydrogel (Fig. 3B), providing a supportive environment where B and T cells reside and interact, mimicking the basic microarchitecture of SLOs.

**Figure 3.**
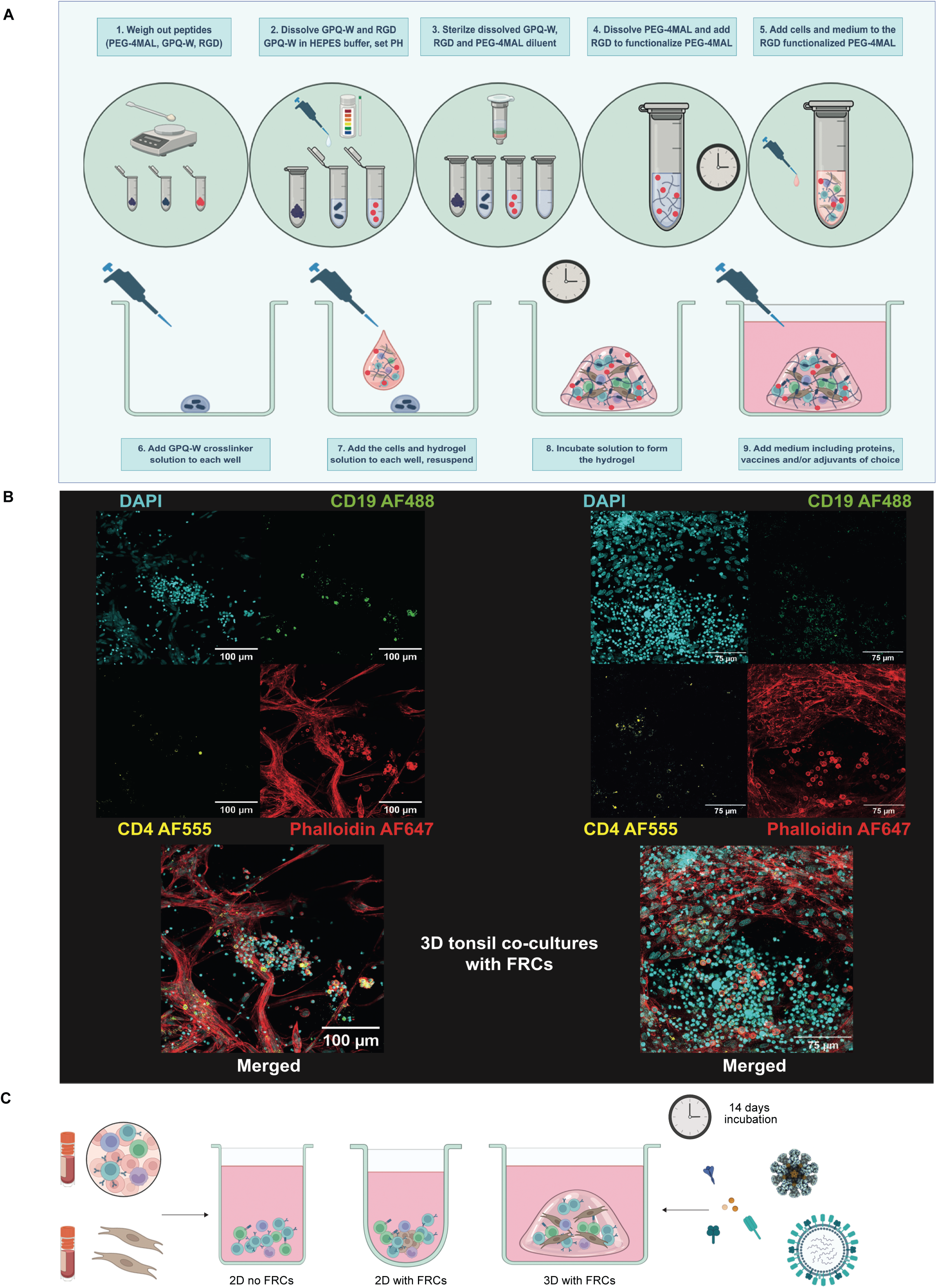
Preparation and confocal imaging of 2D and 3D tonsil co-cultures with and without FRCs. **(A)** Schematic overview of the method used to prepare 3D tonsil co-cultures using a 5% (w/v) 20 kDa PEG-4MAL, RGD functionalized (GRGDSPC), GPQ-W crosslinked (GCRDGPQGIWGQDRCG) hydrogel. The PEG-4MAL macromer, adhesive RGD peptide, and cross-linker GPQ-W were weighed out, dissolved, PH adjusted, and filtered. The PEG-4MAL and RGD solution were combined into a functionalized PEG-4MAL precursor solution. Cells were added to the precursor solution and then mixed with the cross-linking peptide solution pipetted in the bottom of each well. After casting and cross-linking the hydrogel, medium was added. **(B)** Confocal imaging of 3D tonsil co-cultures with FRCs. Cyan = DAPI, green = CD19 Alexa fluor 488, yellow = CD4 Alexa fluor 555, red = phalloidin alexa fluor 647. Left image: influenza vaccine 2022/2023 (Influvac Tetra 2022/2023) stimulated culture on day 7. Scale bars represent 100 μm. Right image: influenza vaccine 2022/2023 stimulated culture on day 14. Scale bars represent 75 μm. **(C)** Cryopreserved tonsil cells and FRCs were thawed and plated out in the two 2D (left and middle) and 3D (right) tonsil (co-)culture setups, after which they were studied for their B and T cell responses for up to 14 days using varying antigen exposures.

Traditional 2D lymphoid cultures were compared to 2D FRC-supported cultures and 3D FRC-supported cultures (Fig. 3C). Medium optimization was carried out in 2D cultures, which showed the lowest overall baseline cell survival. Various types of medium, with and without cytokines, were tested to determine the optimal baseline culture conditions for both antigen-and non-antigen-specific cells in all culture set-ups (Fig. S2A-D). Accordingly, 3D cultures were de-crosslinked to recover cells for flow cytometric evaluation. 2D cultures showed low survival of B cells without FRCs (Fig. S2A), which was not improved through the addition of cytokines IL4 and IL21 (Fig. S2B), but instead caused extensive expansion in the CD8^+^ T cell compartment (Fig. S2C). Interestingly, when FRCs were added the effect of IL4 and IL21 on CD8^+^ T cells was dampened. Even in the absence of stimulation, FRCs significantly improved lymphocyte survival in 2D cultures (P < 0.0001), with further improvements observed when FRCs were combined with a 3D matrix (P < 0.0001; Fig. S2D). This effect was independent of recombinant human B cell activating factor (BAFF) supplementation (Fig. S2D), which was therefore excluded from subsequent cultures. IMDM alone, without additional cytokines, was sufficient to support optimal lymphocyte survival across all culture conditions and was thus selected for further experiments.

### FRC-supported cultures improve lymphocyte viability, differentiation and antibody production after antigen stimulation

Responses of 2D cultures, 2D FRC-supported cultures, and 3D FRC-supported cultures were evaluated when stimulated with various antigen formats (Fig. 3C). These included SARS-CoV-2 WT S or influenza H1N1 HA recombinant proteins, both with and without the adjuvant R848 (TLR7/8-agonist), SARS-CoV-2 WT S nanoparticles (NPs), and two types of influenza vaccines. This setup aimed to mimic conditions in SLOs, where GCs form in response to antigen or vaccine stimulation.

Influenza vaccines induced the most consistent donor-wide responses and are therefore highlighted (Fig. 4); results for the other antigens are presented in Fig. S3. For the CD19^+^ B cell analyses, and for the overall analyses of total antibodies and total T cells, R848-adjuvanted conditions were excluded because this stimulus induced strong non-specific activation (Fig. S3), overpowering antigen specific B and T cell responses with broad overall cellular activation independent of antigen recognition. At day 14, cultures with FRC support contained significantly more viable CD19⁺ B cells than 2D lymphoid cultures without FRCs (2D with FRCs, p = 0.003; 3D with FRCs, p = 0.001), with the highest cell counts in the 3D condition (Fig. 4A, Fig. S3A; Fig. S4). For the CD19^+^ B cell analyses, and for the overall analyses of total antibodies and total T cells, R848-adjuvanted conditions were excluded because this stimulus induces strong non-specific activation (Fig. S3).

**Figure 4.**
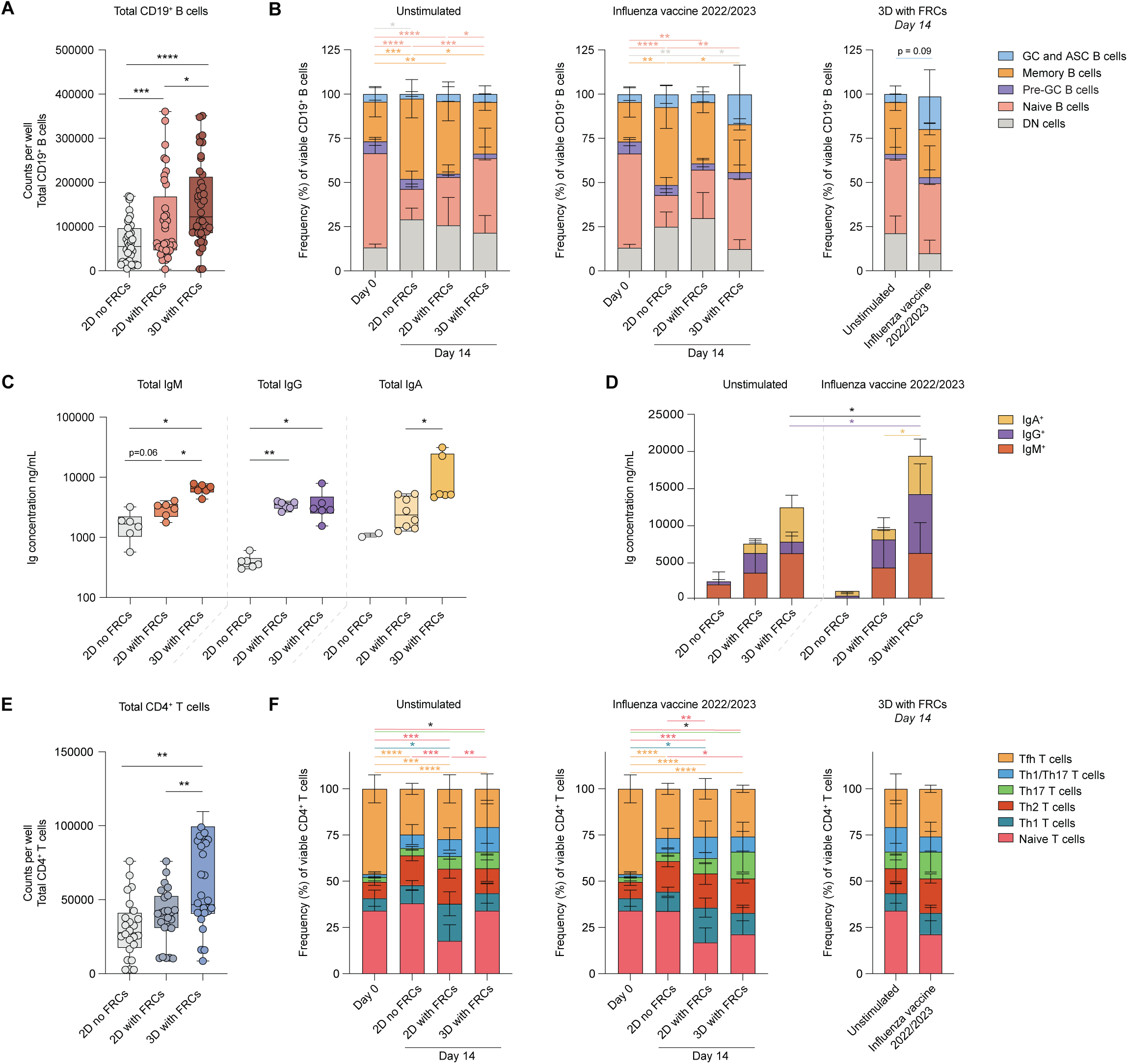
Impact of FRCs and 3D culture on B and T cell survival, differentiation, and immunoglobulin production in response to viral antigens. Survival of B and T cells when cultured in 2D without FRCs, 2D with autologous FRCs, or 3D with autologous FRCs, either left unstimulated or stimulated with influenza vaccine 2022/2023 (Influvac Tetra 2022/2023) **(A)** Total counts of viable CD19⁺ B cells per well at day 14, regardless of stimulation (excluding R848-adjuvanted conditions). **(B)** Percentage of CD19+ B cell subsets, including double-negative cells (DN; CD27^-^CD38^-^IgD^-^), naive B cells (CD27^-^CD38^-^IgD^+^), memory B cells (CD27^+^CD38^-^), pre-GC (CD27^-^CD38^+^), and GC and antibody-secreting cells (GC and ASC; CD27^+^CD38^+^). The left panel shows unstimulated conditions, the middle panel influenza vaccine–stimulated conditions, and the right panel a direct comparison of unstimulated versus influenza vaccine in 3D FRC-supported cultures. **(C)** Concentrations of total IgM, IgG, and IgA (from left to right) in available supernatants collected at day 14, quantified by ELISA across all stimulations (excluding R848-adjuvanted conditions). Ig concentration is in ng/mL. **(D)** Concentrations of IgM, IgG, and IgA in supernatants after 14 days, either unstimulated or stimulated with influenza vaccine. **(E)** Total counts of viable CD4⁺ T cells per well at day 14, regardless of stimulation (excluding R848-adjuvanted conditions). **(F)** Frequency of naive CD4^+^ T cells (CD45RA^+^), Th1 T cells (CD4^+^CD45RA^-^CXCR5^-^CXCR3^+^CCR6^-^), Th2 T cells (CD4^+^CD45RA^-^CXCR5^-^CXCR3^-^CCR6^-^), Th17 T cells (CD4^+^CD45RA^-^CXCR5^-^CXCR3^-^CCR6^+^), Th1/17 T cells (CD4^+^CD45RA^-^CXCR5^-^CXCR3^+^CCR6^+^) and Tfh T cells (CD4^+^CD45RA^-^CXCR5^+^). Baseline (day 0) composition is compared with day 14 cultures. The left panel shows unstimulated conditions, the middle panel influenza vaccine–stimulated conditions, and the right panel a direct comparison of unstimulated versus influenza vaccine in 3D FRC-supported cultures at day 14. Panels A, C, and E were analyzed using paired non-parametric tests (Wilcoxon signed-rank). Panels B, D, and F were analyzed using paired tests with correction for multiple comparisons (Benjamini–Hochberg). Data represent mean ± SD of n = 5 tonsil donors. *=p<0.05, **=p<0.01, ***=p<0.001, ****=p<0.0001; statistical significance is indicated in the color of the corresponding experimental group with a black line depicting differences between total amounts.

Culturing alone reshaped B cell composition over 14 days, shifting from high frequencies of naïve cells to higher proportions of other subsets (Fig. 4B; Fig. S3C). The 3D cultures with FRCs contained the largest naive compartment at day 14, indicating that this condition supports naïve B cell survival (Fig. S3C). In addition, GC and ASC populations were more prominent in 3D FRC-supported cultures than in 2D cultures, particularly after stimulation with the influenza vaccine, which showed a trend toward higher GC/ASC frequencies compared to unstimulated conditions (4-fold increase; p = 0.09; Fig. 4B). In contrast, stimulation with the general activator R848 produced a distinct B cell composition, with even higher GC/ASC frequencies in 3D FRC cultures compared to unstimulated conditions (p = 0.03; Fig. S3C).

Consistent with enhanced B cell survival and differentiation, antibody production was higher in FRC-supported cultures, especially in 3D (Fig. 4C–D). IgM secretion was significantly higher in 3D FRC cultures than in either 2D condition (p = 0.03 for both), with 2D cultures showing only a trend toward higher levels in the presence of FRCs compared to those without (p = 0.06; Fig. 4C). IgG levels were elevated in both FRC-supported conditions compared to 2D cultures without FRCs (p = 0.03 for both; Fig. 4C), while IgA secretion was specifically enhanced in 3D FRC cultures compared to 2D FRCs (p = 0.03). Following influenza vaccine stimulation, antibody secretion increased further in 3D FRC cultures compared to their unstimulated condition (p = 0.02), an effect largely driven by higher IgG levels (p = 0.03; Fig. 4D). R848 stimulation, irrespective of whether combined with spike or HA, strongly enhanced antibody production (Fig. S3D). In 3D FRC-supported cultures, R848 significantly increased IgM, IgG, and IgA secretion (all p < 0.0001), while in 2D FRC cultures it selectively boosted IgM levels (p < 0.0001).

T cell analysis revealed that FRC support improved CD4⁺ T cell recovery as well, with significantly higher counts in 3D cultures with FRCs compared to 2D cultures without FRCs at day 14 (p = 0.003; Fig. 4E; Fig. S3B). Culturing reshaped CD4⁺ T cell composition, with a significant decrease in naive T cells in FRC-supported conditions stimulated with influenza vaccine (2D with FRCs, p = 0.002; 3D with FRCs, p = 0.03; Fig. 4F), accompanied by a relative increase in effector subsets including Th1, Th2, Th17, and Th1/17 (Fig 4F).

Together, these results show that 3D FRC co-cultures provide a supportive niche for both lymphocyte survival and differentiation.

### 3D FRC-supported cultures show amplified specific B cell responses to varying antigen stimulations, with less bystander activation compared to 2D cultures

More detailed analyses were performed on individual antigen-specific B cell responses of each donor, comparing the three tested culture models. Overall, low numbers of antigen-specific B cells were detected in the 2D cultures without FRCs, aligning with the observed overall low survival and activation of B cells (Fig 5A-B, Fig. S5A-B). Despite the higher number of GC/ASC B cells in conditions adjuvanted with R848, there was no preferable increase in the frequency of S- or HA-specific B cells, nor in S- and HA-specific IgGs in the supernatants of these 2D cultures without FRCs (Fig. 5A-D; Fig. S5A-B, E-F). In contrast, 2D FRC-supported cultures showed minor specific responses, mainly in the R848 adjuvanted antigen conditions. In the 3D FRC-supported cultures, a general trend was observed where the percentage of specific B cells was higher across a broader range of tested antigen formats. In a high responsive donor (donor 6; Fig. 5A-D) clear S-specific responses were observed after stimulation with antigen only, as well as R848 adjuvanted antigen, S-NPs and influenza virus and vaccines (Fig. 5A-B), with corresponding increases in S-specific IgM (Fig. S5D) and IgG production (Fig. 5C). Similarly, HA-specific responses were most pronounced following stimulation with influenza HA with R848, the 2022/2023 influenza vaccine, and inactivated influenza virus, showing superior B cell activation across more diverse antigens than in 2D cultures, both at the cellular and antibody levels (Fig. 5B-D). Furthermore, there was an increase in secreted specific antibodies from day 7 to day 14, with the most pronounced responses in the 3D FRC-supported cultures, also including notable increases among lower responding donors (Fig. 5E).

**Figure 5.**
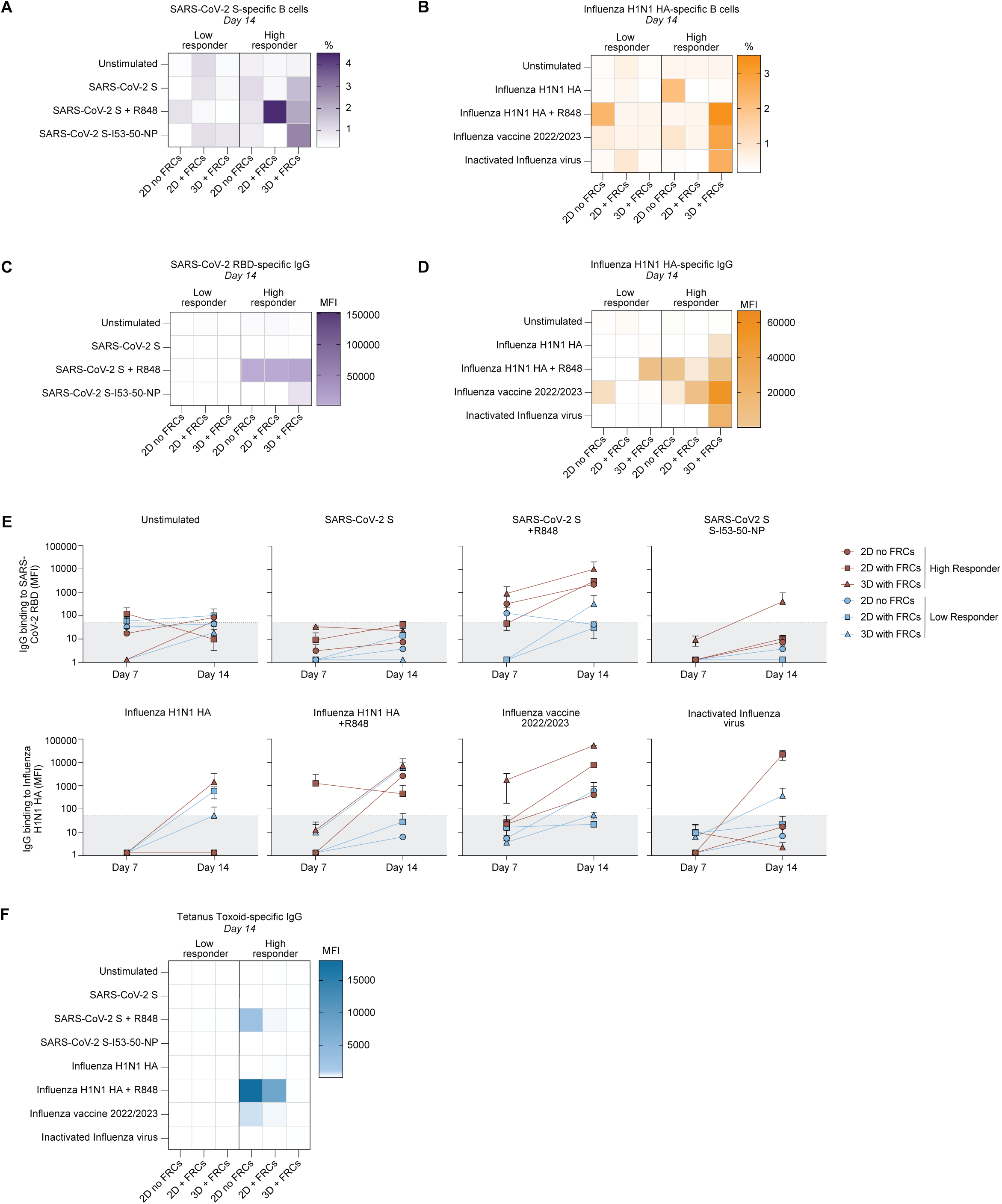
Characterization of influenza and SARS-CoV-2-specific B cell responses, including analysis of antibody production, and bystander effects. Characterization of both Influenza H1N1 hemagglutinin (HA)- and SARS-CoV-2 spike-specific B cell responses after culture. Percentages of protein-specific cells are analyzed using flow cytometry. Protein-specific antibody responses and bystander activation were quantified using a Luminex assay. **(A)** The percentage (%) of SARS-CoV-2 WT spike-specific B cells out of total living CD19^+^ B cells on day 14 for a low-responding tonsil donor (left, tonsil donor 8) and a high-responding tonsil donor (right, tonsil donor 6). The percentage of spike-specific B cells was quantified in an unstimulated culture, or cultures stimulated with either SARS-CoV-2 WT spike (with and without R848) or SARS-CoV-2 WT-I53-50 NPs. All cultures were performed in 2D without FRCs, 2D with autologous FRCs, and 3D with autologous FRCs. **(B)** The percentage (%) of influenza H1N1 HA-specific B cells out of total living CD19^+^ B cells on day 14 for a low-responding tonsil donor (left, tonsil donor 8) and a high-responding tonsil donor (right, tonsil donor 6). The percentage of influenza HA-specific B cells was quantified in an unstimulated culture, or cultures stimulated with either influenza H1N1 HA (with and without R848), influenza vaccine 2022/2023 (Influvac Tetra 2022/2023) or whole inactivated recombinant A/Netherlands/602/2009 influenza virus. All cultures were performed either in 2D without FRCs, 2D with autologous FRCs, and 3D with autologous FRCs. **(C)** SARS-CoV-2 WT receptor-binding domain (RBD)-specific IgG production (mean fluorescence intensity (MFI) minus blank) in the supernatants of the cultures matching the cellular data shown and described in (A). **(D)** Influenza H1N1 HA-specific IgG production (mean fluorescence intensity (MFI) minus blank) in the supernatants of the cultures matching the cellular data shown and described in (B). **(E)** SARS-CoV-2 WT RBD-specific IgG production (MFI minus blank, top row) and influenza H1N1 HA-specific IgG production (MFI minus blank, bottom row) on day 7 versus day 14, either unstimulated or for the varying tested antigen conditions, comparing 2D without FRCs, 2D with autologous FRCs and 3D with autologous FRCs for both the low- and high-responding tonsil donor. **(F)** Tetanus toxoid-specific IgG production (mean fluorescence intensity (MFI minus blank)) in the supernatants of the cultures matching the data shown and described in (A-D). (A-D; F) Data showing the mean of technical duplicates (n=2), (E) data showing the mean ± SD of technical duplicates (n=2).

Alongside quantifying increases in antigen-specific B cells and antibodies, bystander activation was evaluated to assess non-targeted immune responses. This included measuring S-specific antibody production in HA-stimulated conditions and HA-specific antibody production in S-stimulated conditions. Furthermore, antibodies specific to an unrelated antigen, tetanus toxoid, were quantified. An increase in tetanus toxoid-specific IgGs was primarily observed in the R848-adjuvanted and influenza vaccine 22/23 conditions, which generally are the conditions inducing the highest response. Interestingly, this tetanus toxoid-specific bystander activation was mainly seen in the 2D cultures with and without FRCs, but not in the 3D FRC-supported cultures (Fig. 5F). Bystander activation leading to S-specific antibodies in HA-stimulated cultures, and HA-specific antibodies in S-stimulated cultures, were mainly observed in cultures adjuvanted with R848, regardless of the culture set-up used. Interestingly, influenza vaccine stimulated cultures that also contain a broader range of immune activating substances, did not induce these bystander activations as observed in the R848 cultures (Fig. S5G-J).

### 3D FRC-supported cultures enhance antigen-induced ASC differentiation and germinal center-associated chemokine receptor expression

The antigen-induced CD38^+^ B cell responses were further analyzed across the various culture conditions, separating CD38^+^CD20^+^ GC B cells from CD38^++^CD20^-^ ASCs (Fig. S6A). Additional intracellular staining on the lymphocytic fraction at baseline confirmed the CD38^+^CD20^+^ gated GC B cells as a BCL-6 positive B cell population (Fig. S6C).

In 2D cultures without FRCs, GC B cells were detectable and increased significantly after R848 stimulation (p < 0.0001), but ASCs did not develop under any condition (Fig. S6B). Addition of FRCs enabled ASC formation in 2D cultures, although this was largely confined to high-responder donors and most evident with R848 stimulation (Fig. S6B). In contrast, 3D FRC-supported cultures consistently supported differentiation into both GC and ASC compartments across antigen formats, including S protein, S-NPs, and HA (Fig. S6B). Influenza vaccine stimulation reflected the broader donor-dependent pattern observed across antigens (Fig. S6B). In high responders, ASC counts were significantly higher in 3D FRC cultures compared to both 2D conditions (p = 0.0002 vs 2D without FRCs; p = 0.0014 vs 2D with FRCs), and 2D with FRCs also exceeded 2D without FRCs (p = 0.0424; Fig. 6A). In low responders, 3D cultures had significantly higher GC counts than 2D cultures without FRCs (p = 0.005; Fig. 6A). Together, these results highlight the capacity of the 3D stromal microenvironment to drive robust B-cell differentiation into GC and ASC populations, with donor responsiveness shaping the balance between the two compartments.

**Figure 6.**
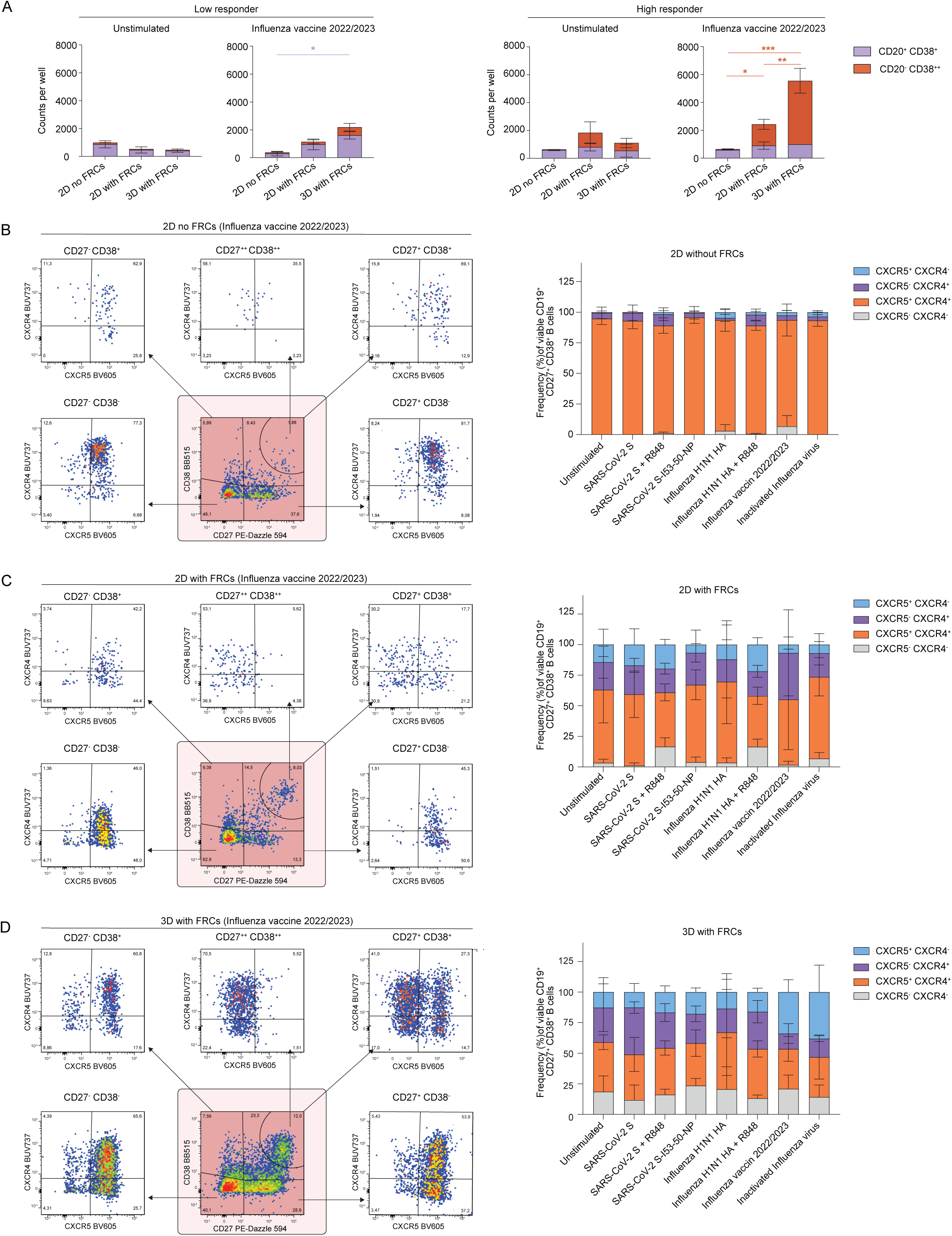
FRCs and 3D culture modulate B cell differentiation and surface chemokine expression in response to viral antigens. B cell differentiation and surface chemokine expression of B cells cultured in 2D without FRCs, 2D with autologous FRCs, or 3D with autologous FRCs, either left unstimulated or stimulated with SARS-CoV-2 WT spike (with and without R848), SARS-CoV-2 WT-I53-50 NP, influenza H1N1 HA (with and without R848), influenza vaccine 2022/2023 (Influvac Tetra 2022/2023) or whole inactivated recombinant A/Netherlands/602/2009 influenza virus. **(A)** Total counts of viable CD19^+^CD20^+^CD38^+^ B cells (GC B cells) and total viable CD19^+^CD20^-^CD38^++^ B cells (ASCs) in a low responder (left; tonsil donor 8) and high responder (right; tonsil donor 6) donor, either unstimulated or stimulated with influenza vaccine 2022/2023 **(B-D)** left: CXCR4 and CXCR5 chemokine expression of total living CD19^+^ B cells, subgated on either naive B cells (CD27^-^CD38^-^), memory B cells (CD27^+^CD38^-^), pre-GC (CD27^-^CD38^+^), GC B cells (CD27^+^CD38^+^) or antibody-secreting cells (CD27^++^CD38^++^), of the high-responding tonsil donor, cultured in 2D without FRCs (B), 2D with autologous FRCs (C), or 3D with autologous FRCs (D), and stimulated with influenza vaccine 2022/2023. Right: overview of the average CXCR4 and CXCR5 chemokine expression of antibody-secreting cells (CD27^+^CD38^+^), cultured in 2D without FRCs (B), 2D with autologous FRCs (C), or 3D with autologous FRCs (D), either unstimulated or stimulated with the varying tested antigen conditions. (A) Data showing the mean ± SD of technical duplicates (n=2), (B-D) data in right panels showing the mean ± SD (n=5 tonsil donors). *=p<0.05, **=p<0.01, ***=p<0.001, ****=p<0.0001, statistical significance is displayed in the colour of the corresponding analyzed experimental group.

Analysis of the GC-associated chemokine receptors CXCR4 and CXCR5 on B cells revealed striking differences between the different culture set-ups. In the 2D cultures without FRCs, B cells mainly co-express both CXCR4 and CXCR5, with no differences for the different B cell populations identified (Fig. 6B, Fig S6D). Both 2D and 3D FRC-supported cultures contained B cell populations expressing either CXCR5 or CXCR4 alone, indicating difference in compartmentalization in the cultures (Fig. S6D). This was also evident in the different B cell subsets across stimuli, with significantly more CXCR5 single-positive B cells among naive (CD27^-^ CD38^-^) and memory B cells (CD27^+^CD38^-^), while CXCR4 single-positive B cells were significantly enriched in the CD27^+^CD38^+^ compartment (2D + FRC vs 2D, *p* < 0.0001; 3D + FRC vs 2D, *p* < 0.0001; Fig. 6C-D). Notably, the CD27^++^CD38^++^ population predominantly consisted of CXCR4 single-positive B cells. This is further reflected by CD38^++^CD20^-^ ASCs, which were primarily CXCR4 single-positive, while CD38^+^CD20^+^ GC B cells contained more CXCR5 and CXCR4 double-positive and CXCR5 single-positive cells, indicative of DZ and LZ GC B cells, respectively (Fig. S6D). While 2D FRC-supported cultures revealed no differences in chemokine receptor response comparing the varying antigen formats, the 3D FRC-supported cultures resulted in more CXCR5 single-positive CD27^+^CD38^+^ B cells when stimulated with both influenza vaccines (Fig. 6D). This increased antigen-induced CXCR5 upregulation and CXCR4 downregulation is mainly found on CD38^++^CD20^-^ ASCs in the 3D model, and less for CD38^+^CD20^+^ GC B cells (Fig. S6D).

### Allogeneic 3D FRC cultures support antigen-specific responses without alloreactive activation

After establishing the beneficial role of FRCs in co-cultures and acknowledging the time-intensive expansion required for autologous FRCs, the use of allogeneic FRCs was explored as an alternative, and their potential was evaluated in side-by-side 3D co-cultures. To exclude potential alloreactive responses induced by allogeneic stromal cells, we first assessed T cell activation markers. Expression of OX40 and CD69 on CD4⁺CD45RA⁻ T cells did not differ between autologous and allogeneic FRC co-cultures at day 7 or 14, and CD8⁺ T cell frequencies remained unchanged, indicating no detectable alloreactive expansion (Fig. 7A–B, Fig. S7A-B). Across stimulations, autologous and allogeneic FRCs also supported comparable T cell viability and differentiation, including GC Tfh frequencies (Fig. 7B, Fig. S7B).

**Figure 7.**
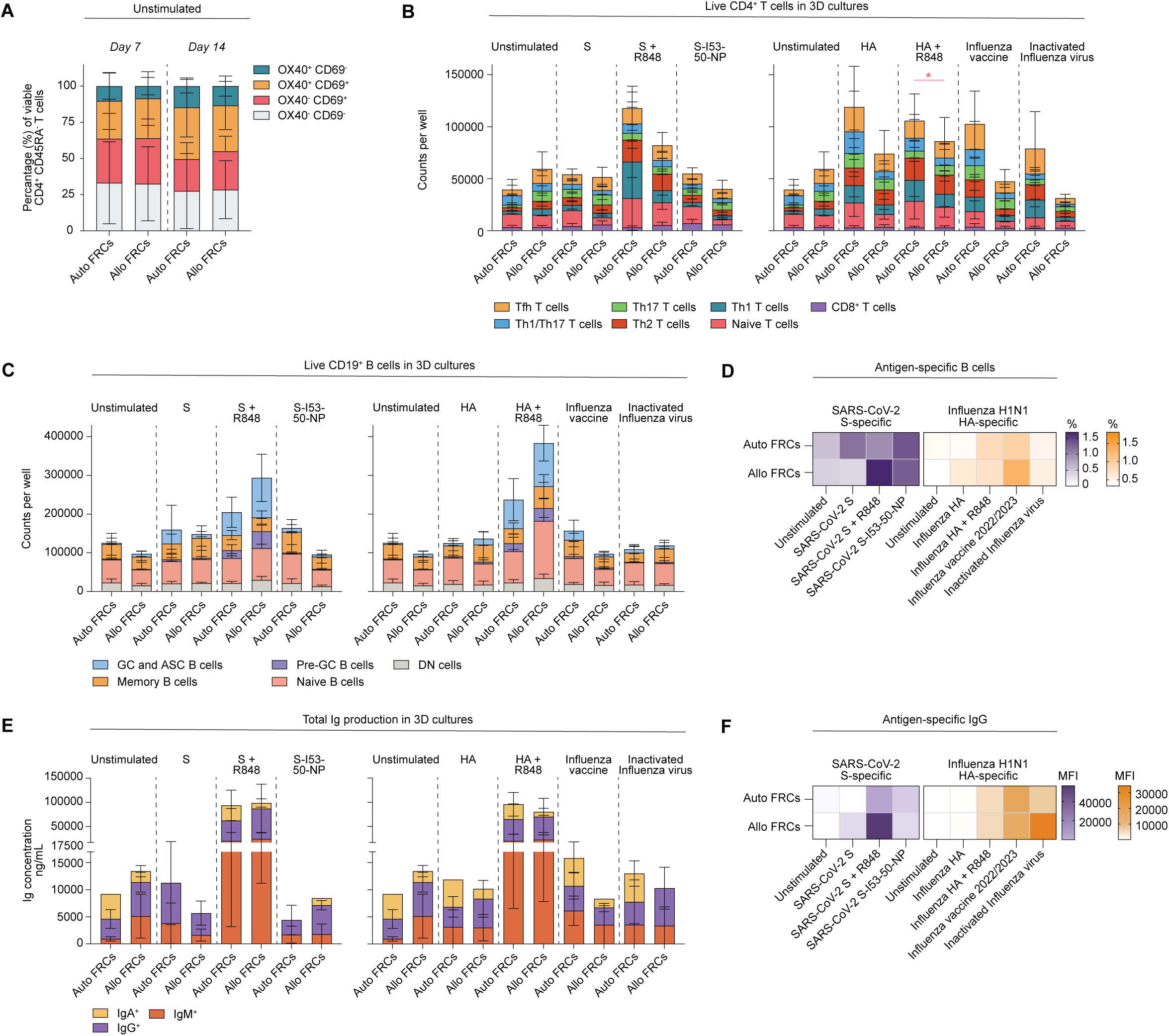
Impact of autologous versus allogeneic FRCs on tonsil cell responses in 3D co-cultures. **(A)** The percentage (%) of viable CD4^+^CD45RA^-^ T cells expressing OX40 and CD69 on day 7 (left) and day 14 (right). Co-cultures were either 2D or 3D with autologous or allogeneic FRCs and were not stimulated with any antigen. **(B)** Total counts of CD8^+^ T cells, naive CD4^+^ T cells (CD45RA^+^), Th1 T cells (CD4^+^CD45RA^-^CXCR5^-^CXCR3^+^CCR6^-^), Th2 T cells (CD4^+^CD45RA^-^ CXCR5^-^CXCR3^-^CCR6^-^), Th17 T cells (CD4^+^CD45RA^-^CXCR5^-^CXCR3^-^CCR6^+^), Th1/17 T cells (CD4^+^CD45RA^-^CXCR5^-^CXCR3^+^CCR6^+^) and Tfh T cells (CD4^+^CD45RA^-^CXCR5^+^) comparing both 2D and 3D autologous and allogeneic co-cultures, cultured unstimulated, or for the varying tested antigen conditions at day 14. **(C)**Total counts of double-negative cells (DN; CD27^-^CD38^-^ IgD^-^), naive B cells (CD27^-^CD38^-^IgD^+^), memory B cells (CD27^+^CD38^-^), pre-GC (CD27^-^CD38^+^), and GC and antibody-secreting cells (GC and ASC; CD27^+^CD38^+^), comparing both 2D and 3D autologous and allogeneic co-cultures, cultured unstimulated, or for the varying tested antigen conditions at day 14. **(D)**The percentage (%) of SARS-CoV-2 WT spike-specific B cells (left) and influenza H1N1 HA-specific B cells (right) out of total living CD19+ B cells on day 14. Co-cultures were either 2D or 3D with autologous or allogeneic FRCs and were either unstimulated or stimulated with the indicated antigen conditions. **(E)** Total immunoglobulin (Ig) production (IgM, IgG, and IgA), quantified in the supernatant either 2D or 3D autologous and allogeneic co-cultures, cultured unstimulated, or for the varying tested antigen conditions. Ig concentration is in ng/mL and quantified at day 14. **(F)** SARS-CoV-2 WT RBD-specific IgG production (left) and influenza H1N1 HA-specific IgG production (right) measured by Luminex assay in supernatants collected on day 14. Co-cultures were either 2D or 3D with autologous or allogeneic FRCs and were either unstimulated or stimulated with the indicated antigen conditions. MFI values are shown after subtracting the background signal (blank). Data showing the mean ± SD (n=5 tonsil donors).

For B cells, viability and differentiation were also similar in autologous and allogeneic FRC co-cultures across unstimulated and antigen-only conditions (Fig. 7A, Fig. S7C). In allogeneic cultures, stimulation with HA and R848 increased naïve B cell viability (p = 0.027), while other B cell compartments remained unaffected (Fig. 7C–D, Fig. S7E–F). Antibody production was comparable between autologous and allogeneic FRC co-cultures (Fig. 7E-F, Fig. S7E-F). Overall, no reproducible differences were detected in B cell responses between autologous and allogeneic FRC conditions.

Taken together, these findings show that allogeneic FRCs do not induce detectable alloreactivity and have minimal effects on B and T cell responses, supporting their use as a practical alternative to autologous FRCs in 3D co-culture systems.

## Discussion

In this study, we developed a 3D FRC-supported *in vitro* model that better mimics human GC-like responses compared to conventional 2D cultures. Our findings demonstrate that the inclusion of FRCs, particularly in a 3D matrix, enhances the survival, differentiation, and activation of B and T cells compared to conventional 2D cultures. By providing structural support within a biomimetic 3D environment, FRCs facilitated the formation of follicle-like structures and promoted GC-like B cell responses. This is evidenced by differential chemokine receptor expression and increased frequencies of antigen-specific B cells and differentiation to ASCs. This enhanced model not only supports antigen-specific antibody responses, but also reduces non-specific, bystander activation, suggesting that 3D FRC-supported cultures may serve as a valuable tool for studying human immune responses and evaluating vaccine candidates.

A 3D environment is fundamental for mimicking GC structure and function, as it better supports cell-cell interactions compared to 2D cultures. Several studies highlighted the benefits of 3D over 2D cultures, demonstrating increased B cell survival GC B cell formation and class-switching (Braham et al. 2023; Zhong et al. 2024; Kramer et al. 2022; Graney et al. 2022; Purwada et al. 2015). This study confirms that 3D cultures improve B and T cell survival and differentiation compared to 2D (FRC-supported), reinforcing the importance of a structured microenvironment for the development of adaptive immune responses.

However, 3D structure alone was insufficient to mimic GC organization. Without FRCs, 3D hydrogels were unstable and disintegrated before the culture period ended, and B and T cells failed to migrate or interact effectively, likely due to the absence of stromal-derived guidance cues. In contrast, FRC-supported 3D cultures facilitated the formation of well-defined follicle-like structures, resembling native GC organization. This structural organization is crucial, as follicle-like aggregates promote immune cell compartmentalization, controlled antigen presentation, and efficient B-T cell interactions, all of which are key for GC-like immune responses (Cremasco et al. 2014; Bajénoff et al. 2006; Panocha et al. 2025).

While previous studies have incorporated stromal support in 3D cultures, these models primarily used mouse fibroblasts overexpressing human CD40L to mimic T cell help rather than replicating the GC stromal compartment (Braham et al. 2023; Zhong et al. 2024; Kramer et al. 2022; Graney et al. 2022; Purwada et al. 2015). As a result, these models lacked the natural B-T cell interactions in an antigen-specific context and did not support migration between GC compartments. Furthermore, many of these models rely on external cytokine supplementation to maintain cell function (Zhong et al. 2024), whereas in our system, immune cells receive cytokine support from other cells in the culture, closely mimicking natural immune cell interaction in SLOs.

FRCs are not only structural components of SLOs but also active regulators of adaptive immune responses. In our 3D culture system, FRCs significantly enhanced antigen-specific B cell frequencies and differentiation into ASCs, as well as GC-associated chemokine receptor expression highlighting their role in B cell selection and maturation. Compared to 2D FRC-supported cultures, 3D cultures displayed a combined response of CD38^+^CD20^+^ GC B cells and CD38^++^CD20^-^ ASCs upon antigen stimulation, resulting in a significantly stronger ASC response. This aligns with previous findings that FRCs help regulate B cell positioning and influence Tfh-B cell interactions (Acton et al. 2021; Wang et al. 2011). These findings underscore that stromal support and a 3D microenvironment together create a more physiologically relevant SLO-like context that supports GC-like organization, antigen-specific immune responses, and B cell differentiation.

Importantly, our data show that FRC-supported 3D cultures selectively enhance antigen-specific responses while minimizing non-specific activation, a factor often overlooked in previous models. While other studies have reported increased antigen-specific immune responses in their lymphoid reaggregated models, they do not distinguish whether these responses were antigen-specific or due to general activation (Wagar et al. 2021; Kastenschmidt et al. 2023; Bonaiti et al. 2024). In contrast, the 2D model, representative of many conventional 2D systems, exhibits extensive bystander activation, highlighting the importance of stromal support in refining immune responses. FRCs are known to regulate immune responses by modulating dendritic cell activity, inducing T cell quiescence, and supporting regulatory T cells (Tregs), all of which contribute to immune tolerance and controlled activation (Link et al. 2007; De Martin et al. 2024; Yu et al. 2017). Combined with the compartmentalization provided by the 3D matrix, these effects may have contributed to the improved antigen specificity.

A defining feature of GCs is their compartmentalization into distinct dark and light zones, which regulate B cell selection and affinity maturation through spatially coordinated B-T cell interactions. This organization gives rise to two distinct GC B cell populations: light zone (CXCR4^lo^CXCR5^+^ or CXCR4^lo^CD83^hi^CD86^hi^) and dark zone GC B cells (CXCR4^hi^CXCR5^+^ or CXCR4^hi^CD83^lo^CD86^lo^) (Allen et al. 2004; Victora et al. 2010, 2012). In this study, CXCR4 and CXCR5 expression on B cells were distinctly different in 3D FRCs co-cultures, with CD38^++^CD20^-^ ASCs containing more CXCR4 single-positive cells compared to cultures without FRCs, and CD38^+^CD20^+^ GC B cells containing both CXCR4 and CXCR5 double-positive and CXCR5 single-positive cells. The differential expression of CXCR4 in CD38^+^CD20^+^ GC B cells suggests active B cell cycling between LZ (CXCR5^+^CXCR4^-^) and DZ (CXCR5^+^CXCR4^+^)-like areas within the FRC-supported cultures. Meanwhile, the absence of CXCR5 on ASCs indicates their readiness to exit the SLO. These results demonstrate the model’s application to offer insights in GC biology research.

Beyond structural organization and GC-like B cell cycling, antigen formats play a critical role in shaping immune responses. Many *in vitro* models rely on live-attenuated viruses or vaccines, which are highly immunogenic and can induce broad, non-specific immune activation (Jeger-Madiot et al., 2024; Wagar et al., 2021). In this study, we tested multiple antigen formats with varying levels of immunogenicity, including recombinant proteins (with and without adjuvants), multivalent antigen displays, whole inactivated viruses, and commercial vaccines derived from SARS-CoV-2 and Influenza H1N1 (pdm09), to evaluate the model’s potential as a tool for vaccine and adjuvant testing. Antigen-specific responses varied by donor immune history and antigen format. The highest responses were observed in a donor with high baseline S-specific B cell frequencies, emphasizing the role of pre-existing immunity in recall responses. Among tested formulations, the Influenza vaccine 22/23 induced strong antigen-specific activation with lower non-specific responses, making it the most controlled immunogen. While recombinant proteins alone elicited minimal responses, recall responses in high-responding donors suggest efficient reactivation of antigen-experienced B cells. Extending culture durations may allow for *de novo* responses, broadening the model’s applicability for studying both primary and recall immunity. Using this model, we also observed that stimulation with R848 (a TLR7/8 agonist) triggered strong but largely antigen-independent responses, suggesting that such adjuvants may not be optimal to use when aiming to elicit specificity *in vitro*. These findings underscore the model’s value as a controlled platform for evaluating vaccine candidates and adjuvants. Future studies should further explore the kinetics and conditions that support *de novo* B cell activation and differentiation.

One parallel advancement recently described in the field of 3D SLO modelling are use of perfused bioreactors or chip models, culturing human tonsil tissue explant fragments, or multicellular hydrogel encapsulated tissue mimics under perfused conditions (Bonaiti et al. 2024; Goyal et al. 2022; Mazzaglia et al. 2024; Jeger-Madiot et al. 2024). Clear benefits of such systems have been described compared to static conditions, such as higher cell viability, increased metabolic activity, and more robust and amplified antigen-specific immune responses. Additionally, our findings demonstrate that autologous and allogeneic FRCs perform comparably in supporting immune responses, reinforcing the scalability of this model for broader applications. The combination of a perfused culture platform, with the optimized biomimetic 3D tonsil environment described in this study would be a next step, to also assess the effect of perfusion to optimized 3D static condition.

This study demonstrates that FRC-supported 3D cultures provide a physiologically relevant model for studying human GC-like responses, surpassing conventional 2D systems. By integrating stromal support within a structured 3D microenvironment, this model enhances B and T cell survival, antigen-specific responses, and ASC differentiation, while also supporting chemokine receptor shuttling and the formation of follicle-like structures, both essential for GC organization and function. These findings establish FRC-supported 3D cultures as a valuable tool for vaccine and adjuvant evaluation, as well as fundamental GC biology research.

## Materials and Methods

### Tonsil cell isolation

Human tonsils were collected with approval by the Medical Ethical Committee of the Amsterdam UMC, Amsterdam, in accordance with the Declaration of Helsinki. Tonsils were obtained from tonsillectomies performed at the Onze Lieve Vrouwe Gasthuis hospital (Amsterdam, The Netherlands) as surgically discarded tissue, without accompanying information on donor age, sex, or clinical history. Collection took place between March 2021 and May 2024. Tonsils were processed to obtain cell suspensions as published, mincing tonsils in small pieces, after which these small tissue fragments were pushed through a metal sieve with a glass plunger, collecting lymphocytes and small tissue fragments in PBS (Breeuwsma and Heesters 2023). Lymphocytes were collected from this solution by density gradient centrifugation using Lymphoprep (Axis-Shield) and cryopreserved before use in experiments. The remaining small tissue fragments were collected and plated in Dulbecco’s Modified Eagle Medium high glucose (DMEM high glucose, Gibco) with 10% (v/v) fetal calf serum (FCS; Bodinco), 100 U/mL Penicillin (Life Technologies) and 100 μg/mL Streptomycin (Life technologies). Upon outgrowth of stromal, adherent cells, a round of trypsinization was performed, removing residual tissue fragments using a 100 μm cell strainer (Corning). The resulting fibroblastic reticular cells (FRCs) were cultured for another passage until reaching confluency, and cryopreserved before use in experiments.

### Protein production and purification

The tetanus toxoid protein was acquired from Creative Biolabs. All protein constructs, including soluble SARS-CoV-2 WT spike (pre-fusion stabilized with a T4 trimerization domain (Brouwer et al. 2020)), SARS-CoV-2 WT RBD (van der Straten et al. 2024), Influenza H1N1 HA (H1N1pdm2009, A/Netherlands/602/2009, GenBank: CY039527, (Aartse et al. 2021)), SARS-CoV-2-S-I5350A.1NT1 plasmid (Brouwer et al. 2021), with or without avi-tag, were designed as previously described. The protein production and purification of all constructs were performed as previously described (Brouwer et al. 2020). For protein production, HEK 293F cells (Invitrogen) were maintained in Freestyle medium (Life Technologies) at a density of 0.8-1.2 million cells/mL. Cells were transiently transfected with the respective expression plasmids using polyethylenimine hydrochloride (PEI) MAX (Polysciences). The transfection mixture consisted of PEI MAX at a concentration of 1 mg/mL and expression plasmids at 312.5 µg/L in a 3:1 ratio (PEI MAX:plasmid) in OptiMEM (Gibco).

This mixture was added to the cells and six days after transfection, supernatants were harvested by centrifugation at 4000 rpm for 30 minutes at 4 degrees. Supernatants were filtered using 0.22 μm Steritop filters (Merck Millipore). His-tagged proteins were purified from the clarified supernatant by affinity chromatography using Ni-NTA agarose beads (QIAGEN). Following elution, proteins were concentrated, and buffer exchanged into PBS using Vivaspin centrifugal concentrators with a 100 kDa molecular weight cutoff (MWCO) (GE Healthcare). After purification, Avi-tagged proteins were biotinylated using a BirA500 biotin-ligase reaction kit (Avidity), according to the manufacturer’s instructions. To remove unbound biotin, the proteins were further purified using size-exclusion chromatography (SEC) on a Superdex 200 16/60 column (Cytiva), with PBS as the elution buffer.

For the assembly of SARS-CoV-2 WT Spike I53-50 nanoparticles (NP), the purified SARS-CoV-2-S-I5350A.1NT1 fusion proteins were buffer exchanged into Tris-buffered saline (TBS) and sterilized using a 0.22 μm spin column. NP assembly was performed as previously described (Brouwer et al. 2021). Briefly, the S-I53-50A.1NT1 proteins were further purified by size-exclusion chromatography (SEC) on a Superose 6 increase 10/300 GL column (GE Healthcare) in TBS with 5% glycerol. The appropriate fractions were collected and pooled. Equimolar amounts of I53-50B.4PT1 were mixed with the S-I53-50A.1NT1 protein and incubated overnight to allow NP assembly. Assembled NPs were then purified by SEC to remove any unassembled components, and the relevant fractions were concentrated using a 10.000 Da Vivaspin column (GE healthcare).

Protein concentrations of all proteins were determined using a Nanodrop 2000 Spectrophotometer using the protein’s peptidic molecular weight.

### Vaccines

Influvac Tetra 2022/2023 (Abbott Biologicals BV, the Netherlands) containing inactivated surface antigens (hemagglutinin (HA) and neuraminidase (NA)) of the following virus strains: A/Victoria/2570/2019 IVR-215, A/Darwin/9/2021 SAN-010 B/Austria/1359417/2021, BVR-26 and B/Phuket/3073/2013 wild type (WT), containing 30 μg/mL HA of each strain.

### Whole inactivated recombinant A/Netherlands/602/2009 influenza virus

293T cells (ATCC) were cultured in Dulbecco modified Eagle’s medium (DMEM) (Capricorn Scientific) supplemented with 10% fetal calf serum (FCS) (Sigma-Aldrich),1x non-essential amino acids (Capricorn Scientific), 1 mM sodium pyruvate (Gibco), 2 mM L-glutamine (Capricorn Scientific), 100 U/mL penicillin and 100 U/mL streptomycin (Capricorn Scientific), and 0.5 mg/mL Geneticin (Gibco). Madin-Darby canine kidney (MDCK) cells (ATCC) were cultured in Eagle’s minimal essential medium (EMEM, Capricorn Scientific), supplemented with 10% FCS, 1x non-essential amino acids (Capricorn Scientific), 1.5 mg/mL sodium bicarbonate (Gibco), 10 mM HEPES (Capricorn Scientific), 2 mM L-glutamine (Capricorn Scientific), 100 U/mL penicillin and 100 U/mL streptomycin (Capricorn Scientific). Cells were cultured at 37 °C, 5% CO2, and passaged twice weekly.

The A/Netherlands/602/2009 recombinant influenza virus was generated by reverse genetics using eight bidirectional plasmids as described previously (Siegers et al. 2023). It contained the hemagglutinin and neuraminidase of the A/Netherlands/602/2009 virus and the remaining six genes of A/Puerto-Rico/8/1934. Briefly, recombinant virus was produced by transfecting bidirectional plasmids into 293T cells using the calcium-phosphate transfection method. Approximately 16 hours after transfection, the cells were washed once with PBS and fresh media containing 2% FCS. Three days after transfection, supernatant dilutions from 293T cells were used to inoculate MDCK cells. Virus stock production in MDCK cells was performed using EMEM medium containing the same supplements as described above, but without FCS and with the addition of 20-35 mg/mL of N-tosyl-L-phenylalanine chloromethyl ketone (TPCK)-treated trypsin (Sigma-Aldrich), referred to as infection medium. MDCK supernatant was harvested two to three days post-inoculation and centrifuged at 2,100 g for 10 minutes to remove cellular debris. The presence of virus was confirmed by hemagglutination (HA) assays using 1% turkey red blood cells (TRBCs, harvested from in-house turkeys) in PBS. The sequences of all plasmids, along with the hemagglutinin and neuraminidase genes of A/Netherlands/602/2009 recombinant viruses, were confirmed with Sanger sequencing using the BigDye™ Terminator v3.1 Cycle Sequencing Kit (Applied Biosystems) and the 3500xL Genetic Analyzer (Applied Biosystems).

Whole-inactivated virus vaccines were generated as follows. Eleven-day old embryonated chicken eggs were inoculated with the A/Netherlands/602/2009 recombinant influenza virus. Allantoic fluid was harvested two days post-inoculation and centrifuged for ten minutes at 2,100 g to remove cellular debris. Subsequent centrifugation steps were performed at 124,000 g (SW 32 Ti, Beckman Coulter) at 4 °C, unless indicated otherwise. The allantoic fluid was concentrated on a 60% sucrose cushion by centrifuging for 2 hours. Following this, resuspended sucrose cushions from multiple tubes were pooled and loaded on 60-50-40-30-20% sucrose gradients, which were centrifuged overnight at the lowest deceleration setting. The virus band, located on top of the 30% sucrose layer, was harvested, diluted in PBS, and subsequently pelleted by centrifugation for 2 hours to remove the sucrose. The pellet was dissolved in PBS. The dissolved pellets were transferred to dialysis chambers (Slide-A-Lyzer™ Dialysis Cassettes, 10K MWCO, Thermo Fisher Scientific) which were subsequently submerged in PBS containing 0.01% formalin for three days. The dialysis chambers were then submerged in PBS for a day, during which the PBS was refreshed twice. The resulting vaccines were aliquoted and stored at −80 °C. Vaccine inactivation was confirmed by two serial blind passages on MDCK cells. Total protein content was determined using the Pierce BCA total protein analysis kit (Thermo Fisher Scientific). The absolute HA content was estimated from non-reducing SDS-PAGE protein gels using dilutions of a A/Netherlands/602/2009 hemagglutinin recombinant protein as standard and stained with instant Blue (Expedeon).

### 2D and 3D tonsil co-cultures

FRCs were thawed and expanded before the start of the experiments. Tonsil lymphocytes were thawed on the day of 2D and 3D (co-)culture plating. All conditions were performed using 250.000 tonsil lymphocytes per well. Cultures with FRCs contained 50.000 FRCs per well. 2D cultures without FRCs were performed in 96 Well Sphera Low-Attachment Surface plates (ThermoFisher). 2D cultures with FRCs were performed in non-tissue culture-treated 96-well U bottom plates (Greiner), in which plated FRCs formed spheroids. 3D cultures were performed in non-tissue culture-treated 48-well plates (ThermoFisher). All cultures were performed using B cell medium as a basis: Iscove’s Modified Dulbecco’s Medium (IMDM, Gibco), 10%(v/v) fetal calf serum (FCS, Bodinco), 100 U/ml penicillin (Life Technologies), 100 μg/ml streptomycin (Life Technologies), 2 mM L-glutamine (Life Technologies), 50 μM β-mercaptoethanol (Sigma Aldrich), and 20 μg/ml Transferrin depleted for IgG (Sigma Aldrich). Conditions with antigen stimulation contained either SARS-CoV-2 WT spike protein (1 µg/ml, with and without 1 µg/ml Resiquimod (R848, MedChemExpress), SARS-CoV-2 WT-I53-50 nanoparticles (1.219 µg/ml), influenza H1N1 HA protein (1 µg/ml, with and without 1 µg/ml R848), influenza vaccine 2022/2023 (Influvac Tetra 2022/2023, 1 µg/ml of total HA) or whole inactivated recombinant A/Netherlands/602/2009 influenza virus (0,01 µg/ml of total HA). 3D cultures were performed in 15 µL PEG-4MAL-RGD functionalized hydrogels. All cultures were analyzed on day 0 (input lymphoid cells), 7 days after culture and 14 days after culture.

### PEG-4MAL hydrogel

A cell encapsulating hydrogel was prepared as described (Braham et al. 2023; Cruz-Acuña et al. 2018), using a four-armed polyethylene glycol macromer with maleimide groups at each terminus (PEG-4MAL, MW 20 kDa, Laysan Bio), to which cysteine-containing adhesive peptide RGD (GRGDSPC, AAPPTec, custom synthesis, purity: >95%, trifluoroacetic acid (TFA) removal) was conjugated. The gel was crosslinked using protease degradable cross-linking peptide GPQ-W (GCRDGPQGIWGQDRCG, AAPPTec, custom synthesis, purity: >95%, (TFA) removal). In short: an aliquot of the PEG-4MAL macromer, adhesive RGD peptide, and cross-linker GPQ-W were allowed to reach room temperature. The needed amount of GPQ-W and RGD was weighed out and dissolved using 20 mM HEPES buffer (Sigma), after which the PH was adjusted to 7.4. Both solutions were filtered using a Costar Spin-X centrifuge tube (Corning). The PEG-4MAL macromer was dissolved in sterile, prefiltered 20 mM HEPES buffer. The PEG-4MAL and RGD solution were combined in a 2:1 ratio and incubated for 15 min at 37°C. Cells were resuspended in B cell medium after which the functionalized PEG-4MAL-RGD precursor was added. The cross-linking peptide solution (GPQ-W) was added to the bottom of each well. The functionalized PEG-4MAL-RGD precursor mixture containing the cells was pipetted on top of the cross-linking peptide solution and re-suspended quickly, in a total final volume of 15 µL per well. After mixing the hydrogel, it was left cross-linking at 37°C for 20 min, after which medium was added (illustrated in Fig. 3B).

### Cell collection and decrosslinking of PEG-4MAL hydrogel

#### 2D cultures without FRCs

Cultures were resuspended and moved to a new 96-well V-bottom plate (ThermoFisher Scientific). After spinning down the plate, the supernatant was removed and stored at −20°C. Cells were washed two times with PBS containing 0.1% (w/v) bovine serum albumin (BSA, Sigma Aldrich) and left in PBS containing 0.1% (w/v) BSA until further staining and analysis.

#### 2D cultures with FRCs

Cultures were resuspended and moved to a new 96-well V-bottom plate (ThermoFisher Scientific). After spinning down the plate, the supernatant was removed and stored at −20°C. Cells were washed two times with PBS-0.1% BSA. each time letting the plate stand for 30 seconds, letting FRC spheroids sediment. The single cell suspension above was collected and moved to a new 96-well V-bottom plate, removing these clumps. After washing and removal of the FRCs, cells were left in PBS-0.1% BSA until further staining and analysis.

#### 3D cultures

Supernatant was removed and stored at −20°C. Cells were washed once with PBS. Trypsin was added to the 48-well plate and incubated for 3 minutes. To inactivate the trypsin, B cell medium was added in a 1:1 ratio. The trypsin diluted in B cell medium was transferred to the V-bottom 96-well plate, to collect free-floating cells. Next, 200 U/ml collagenase type 1 (Gibco) diluted in B cell medium was added to each gel in the 48-well plate. Gels were mechanically disrupted using a positive displacement pipette. The decrosslinking solution was incubated for 1 hour at 37°C. All collected cells were transferred to a 96-well V-bottom plate (ThermoFisher Scientific). The cells in the 48-well plate were washed two times with PBS-0.1% BSA and every time the cells were transferred to the V-bottom 96-well plate. Cells of the multiple collection rounds were pooled per well. After collection of all cells, the plate was left standing for 30 seconds, to let residual FRC clumps sediment. The single cell suspension above was collected and moved to a new 96-well V-bottom plate, removing these clumps. After this, the cells were stained for further analysis.

### Probe generation

Biotinylated protein antigens were individually multimerized with Alexa Fluor 647 and BV421 (both Biolegend) labeled streptavidin, as described previously (Brouwer et al. 2020). In short, biotinylated proteins and fluorescently tagged streptavidin were combined in a 2:1 molar ratio of protein to fluorochrome and then incubated for one hour at 4°C. Free streptavidin conjugates were quenched with 10 µM biotin (Genecopoiea) for a minimum of 10 minutes. The individually labeled proteins were then combined in equimolar amounts to achieve a final concentration of 45.5 pM.

### Flow cytometric characterization

Anti-human antibodies used for flow cytometric analysis are:

#### B cell panel

Brilliant Ultraviolet 737 anti-human CXCR4 Antibody (BD Biosciences, Clone: 12G5), Brilliant Violet 605 anti-human CXCR5 Antibody (Biolegend, Clone: RF8B2), Alexa Fluor 700 anti-human IgG Antibody (BD Biosciences, Clone: G18-145), LIVE/DEAD™ Fixable Near IR (780) Viability Kit (Invitrogen, L34992), eFluor780 anti-human CD3 Antibody (Invitrogen, Clone: UCHT1), eFluor780 anti-human CD14 Antibody (Invitrogen, Clone: 61D3), eFluor780 anti-human CD16 Antibody (Invitrogen, Clone: CB16), PE anti-human IgA Antibody (SouthernBiotech), PE-Dazzle 594 anti-human CD27 Antibody (Biolegend, Clone: O323), PE-Cy7 anti-human IgM Antibody (Biolegend, Clone: MHM-88), Brilliant Blue 515 anti-human CD38 Antibody (BD Biosciences, Clone: HIT2), Brilliant Ultraviolet 395 anti-human IgD Antibody (BD Biosciences, Clone: IA6-2), Brilliant Ultraviolet 496 anti-human CD19 Antibody (BD Biosciences, Clone: SJ25C1), Brilliant Ultraviolet 805 anti-human CD20 Antibody (BD Biosciences, Clone: 2H7), and Brilliant Stain Buffer Plus (BD Horizon).

#### T cell panel

PE-Dazzle 594 anti-human CD185 Antibody (CXCR5, Biolegend, Clone: J252D4), Brilliant Blue 515 anti-human CD196 Antibody (CCR6, BD Biosciences, Clone: 11A9), Brilliant Violet 605 anti-human CD183 Antibody (CXCR3, Biolegend, Clone: G025H7), APC anti-human CD279 Antibody (PD-1, Invitrogen, Clone: J105), LIVE/DEAD™ Fixable Near IR (780) Viability Kit (Invitrogen, L34992), PE anti-human CD134 Antibody (OX40, BD Biosciences, Clone: ACT35), PE-Cy7 anti-human CD45RA Antibody (Biolegend, Clone: HI100), Brilliant Violet 421 anti-human CD154 Antibody (CD40L, BD Biosciences, Clone: TRAP1), Brilliant Ultraviolet 395 anti-human CD3 Antibody (BD Biosciences, Clone: HIT3a), Brilliant Ultraviolet 496 anti-human CD4 Antibody (BD Biosciences, Clone: OKT4), Brilliant Ultraviolet 737 anti-human CD69 Antibody (BD Biosciences, Clone: FN50), Brilliant Ultraviolet 805 anti-human CD8 Antibody (BD Biosciences, Clone: SK1), and Brilliant Stain Buffer Plus (BD Horizon).

#### FRC panel

APC anti-human CD45 (Beckman Coulter, Clone: J33), LIVE/DEAD™ Fixable Near IR (780) Viability Kit (Invitrogen, L34992), PE anti-human CD35 (Biolegend, Clone: E11), PE-Cy7 anti-human PDPN (Biolegend, Clone: NC-08), FITC anti-human CD31 (Beckman Coulter, Clone: 5.6E), PerCP-Efluor710 anti-human HLA-ABC (Invitrogen, Clone: W6/32), BV421 anti-human HLA-DR (BD Horizon, Clone: G46-6), BUV395 anti-human CD21 (BD Optibuild, Clone: B-ly4).

#### BCL-6 and c-Myc panel

PerCP-eFluor 710 anti-human CXCR4 Antibody (eBioscience, Clone: 12G5), Brilliant Violet 605 anti-human CXCR5 Antibody (Biolegend, Clone: J252D4), Alexa Fluor 700 anti-human IgG Antibody (BD Biosciences, Clone: G18-145), LIVE/DEAD™ Fixable Near IR (780) Viability Kit (Invitrogen, L34992), PE anti-human IgA Antibody (SouthernBiotech), PE-Dazzle 594 anti-human CD27 Antibody (Biolegend, Clone: O323), Brilliant Blue 515 anti-human CD38 Antibody (BD Biosciences, Clone: HIT2), Brilliant Violet 421 anti-human CD86 Antibody (BD Biosciences, Clone: 2331), Brilliant Violet 510 anti-human IgM Antibody (Biolegend, Clone: MHM-88), Brilliant Ultraviolet 395 anti-human IgD Antibody (BD Biosciences, Clone: IA6-2), Brilliant Ultraviolet 496 anti-human CD19 Antibody (BD Biosciences, Clone: SJ25C1), Brilliant Ultraviolet 737 anti-human CD138 Antibody (BD Horizon, Clone: ML15), Brilliant Ultraviolet 805 anti-human CD20 Antibody (BD Biosciences, Clone: 2H7), APC anti-human c-Myc Antibody (Biolegend, Clone: 9E10), and PE-Cy7 anti-human BCL6 Antibody (Biolegend, Clone: 7D1).

#### Extracellular staining panels

Cells were resuspended in FACS buffer (PBS-0.1% BSA) containing antibodies and, for the panels containing chemokine receptors, incubated first at 37°C with the chemokine receptor antibodies for 30 minutes. Next, all other antibodies were added for subsequent incubation at 4°C for 30 min. After incubation, cells were washed and fixated using 4% paraformaldehyde (PFA, Sigma). Fixated cells were washed and then re-suspended in FACS buffer.

#### Intracellular BCL-6 and c-Myc staining

After extracellular chemokine and antibody mix staining, cells were washed and fixated using Foxp3 fixation buffer (eBioscience). Foxp3 fixed samples were washed once with Foxp3 permeabilization buffer (eBioscience). Samples were stained in 25µL staining mix containing antibodies against BCL6 and c-myc and incubated overnight at 4°C. On the next day, samples were washed once with Foxp3 permeabilization buffer and resuspended in FACS buffer.

Flow cytometric data was acquired using a BD FACSymphony (Becton Dickinson).

### Immunocytochemistry

3D cultures were washed with PBS and then fixated in 4% PFA for 1 hour. Samples were washed with PBS and blocked with 5% (w/v) BSA in PBS for 24 hours at 4°C on a plate shaker (100 1/min) and subsequently washed in PBS. Immunodetection was performed by incubation with purified mouse anti-human CD19 antibody (14.3 ug/mL, HIB19, BD Biosciences) and purified rat anti-human CD4 antibody (14.3 ug/mL, A161A1, Biolegend) for the first panel and by incubation with purified mouse anti-human CD185 (CXCR5) antibody (14.3 ug/mL, J252D4, Biolegend), recombinant rabbit anti-CXCR4 antibody (77.4 ug/mL, UMB2, Abcam) and rat CD19 monoclonal antibody (14.3 ug/mL, 6OMP31, Invitrogen) for the second panel for 72h at 4°C. As a second step, goat anti-mouse IgG (minimal x-reactivity) Alexa Fluor 488 (40 ug/mL, 405319, Biolegend), goat anti-rat IgG (minimal x-reactivity) Alexa Fluor 555 (40 ug/mL, 405420, Biolegend) were added for both panels, with additionally goat anti-rabbit IgG H&L Alexa Fluor® 647 (40 ug/mL, ab150079, Abcam) for the second panel for 72 hour at 4°C, all in PBS-0.1% BSA and incubated on a plate shaker (100 1/min). All antibodies were centrifuged before use (13.000 rpm for 5 min). Between staining steps, samples were washed in 0.1% Tween 20 (v/v, Merck Millipore) in PBS for 2 hours (first wash) and for 24 hours at 4°C on a plate shaker (100 1/min, second wash). Samples were then stained for DAPI (100 ng/mL, D9542, Sigma Aldrich) and Phalloidin Alexa Fluor™ 647 (165 nM, A22287, Invitrogen™) for the first panel, and DAPI and Phalloidin-Atto 700 (0.4 nmol/mL, 79286, Sigma Aldrich) for the second panel for 24 h at 4°C. Then samples were washed twice in PBS, with a final washing step overnight at 4°C on a plate shaker (100 1/min). All samples were mounted on microscopy slides using Vectashield (Vector Laboratories, H-1700-10).

### Confocal Imaging

Confocal images were taken with a Leica TCS SP8 or a Leica Stellaris 8 confocal microscope using Leica LASX acquisition software. On the SP8, four detection channels collected the fluorescent signal from the used fluorochromes: DAPI, Alexa Fluor 488, Alexa Fluor 555 and Alexa Fluor 647, which were given the pseudocolors cyan, green, yellow, and red. On the Stellaris 8, five detection channels collected the fluorescent signal from the used fluorochromes: DAPI, Alexa Fluor 488, Alexa Fluor 555, Alexa Fluor 647, and Phalloidin–Atto 700, which were given the pseudocolors blue, green, yellow, cyan and red.

### ELISA

96-well Nunc Maxisorp plates (ThermoFisher Scientific) were coated with 2 ug/ml monoclonal mouse anti-human IgG (MH16-1, Sanquin), 1 ug/ml monoclonal mouse anti-human IgA (MH14-1, Sanquin) or 2 ug/ml monoclonal mouse anti-human IgM (MH15-1, Sanquin) diluted in PBS and incubated overnight at RT. The plates were washed 5 times with PBS-Tween (10% (v/v)). The supernatants of the cultures were thawed and diluted 1:25, 1:125, 1:625 in High-Performance ELISA buffer (HPE buffer, Sanquin) and added to the plates. A standard curve of a serum pool with known antibody concentrations (Pool serum 15-123, Sanquin) was diluted in HPE and also added to the plate, as well as 2 blanks. The plates were incubated for 1 hour at RT on a shaker and washed afterwards. Then, 1 ug/ml monoclonal mouse anti-human IgG HRP (MH16-1, Sanquin), 1 ug/ml monoclonal mouse anti-human IgA HRP (MH14-1, Sanquin), or 1 ug/ml monoclonal mouse anti-human IgM (MH15-1, Sanquin), all diluted to the final concentrations in HPE, were added to the plate and incubated for 1 hour at RT on a shaker. The plates were washed and the substrate, 1-step Ultra TMB-ELISA (Thermo Fisher Scientific) diluted in MiliQ in a ratio of 1:1, was added to the plate. The reaction was stopped with 0.2 M H_2_SO_4_, measured at 450 nm and corrected for absorbance background at 540 nm with a Synergy 2 microplate reader (Biotek Instruments Inc.).

### Multiplex Immunoassay

The humoral response in the supernatant of tonsil cultures was measured using a customized in-house Luminex immunoassay (Grobben et al. 2021). Briefly, HIV-1 ConM Env v.7, tetanus toxoid, SARS-CoV-2 Wuhan Spike protein, SARS-CoV-2 Wuhan RBD, Influenza HA H1N1 PDMN2009, and a no-protein control (75 μg each, equimolar corrected) were covalently coupled to MagPlex beads (12.5 million beads per protein) using a two-step carbodiimide reaction. Supernatant samples were diluted 1:2 and incubated with the beads overnight, followed by detection with 1,3 ug/ml goat anti-mouse IgM-PE or IgG-PE (Southern Biotech). The mean fluorescence intensity (MFI) was measured using the Magpix platform (Luminex), capturing the median signal from around 50 beads per well. Background fluorescence was accounted for by subtracting the MFI values from buffer and beads-only controls.

### Quantification and statistical analysis

Flow cytometric data was analyzed using FlowJo v10.8.1 software. The gating strategy for both the B cell and T cell panels is shown in the supplements (Supplementary Fig. S1). The cell counts that are presented are corrected for the original total volume the cells were in. IgG, IgA and IgM concentrations were calculated using ELISA-Logit-V01Jul2018. Graphs were made using GraphPad Prism 9.1.1. Fluorescent z-stack images were processed using Fiji 1.53t software to create single maximum projections. Image overviews of the 3D cultures were merged using the mosaic function of the Leica LASX software, stitching the images together using smooth and linear blending.

## Acknowledgements

We would like to thank: Mitch Brinkkemper and Rogier W. Sanders (Department of Medical Microbiology and Infection Prevention, Amsterdam UMC the Netherlands) for their kind sharing of produced spike nanoparticles, Neil P. King (University of Washington, Seattle, USA) for kindly providing the I53-50B.4PT1 protein, Mariel Duurland and Charlotte Menage (both Sanquin, Amsterdam, the Netherlands) for their kind sharing of produced (biotinylated) spike protein, Wouter Olijhoek (Department of Medical Microbiology and Infection Prevention, Amsterdam UMC the Netherlands) for his kind help in isolating FRCs and producing (biotinylated) spike protein, Nienke Haverkate (Laboratory of Experimental Immunology, Amsterdam UMC, the Netherlands) for facilitating the acquisition of tonsillar tissue, the surgeons and patients of the Onze Lieve Vrouwen Gasthuis (Amsterdam, the Netherlands) for their generous contribution of tonsil tissue for this study, Simon Tol and Erik Mul (both Sanquin, Amsterdam, the Netherlands) for their kind help and troubleshooting during our hours of running all samples in the Cell Analysis Facility and Daisy Picavet-Havic (Microscopy Core Facility, Amsterdam UMC, the Netherlands) for her kind help with setting up the confocal microscope.

The illustrations shown in Figures 1 and 3 were created with BioRender.com.

## Author contributions

Conceptualization: MJVG, ATB, MVJB, MDG, CACMVE, JDW, MC, SMVH.

Funding acquisition: MJVG, ATB, CACMVE, JDW, SMVH.

Investigation: MVJB, MDG, LB, SK, TMB

Methodology: MJVG, ATB, MVJB, MDG, CACMVE, JDW, MC, SMVH, MR.

Resources: CACMVE, JDW, SMVH, ATB, MJVG, MR

Supervision: MJVG, ATB, CACMVE, JDW, MC, SMVH.

Writing - original draft: MVJB and MDG

Writing - review and editing: MJVG, ATB, MVJB, MDG, CACMVE, JDW, MC, SMVH, MR.

---

**Figure S1. Flow cytometry gating strategy for tonsil lymphocytes.** Flow cytometry staining and gating strategy, showing representative data of tonsil lymphocytes analyzed for B cells **(A)** and T cells **(B)** on day 0.

**Figure S2. Culture optimization of unstimulated tonsil cells, using varying (co-)culture conditions, basal media, and cytokine supplementation. (A)** Total viable CD19^+^ B cell counts on day 14, comparing 2D cultured tonsil cells without FRCs to 2D cultured tonsil cells with FRCs, either in RPMI versus IMDM, with or without additional supplementation of B cell cytokines IL4 and IL21 (both 50 ng/mL). **(B)** Total counts of naive B cells (CD27^-^CD38^-^), memory B cells (CD27^+^CD38^-^), pre-GC (CD27^-^CD38^+^), and GC and antibody-secreting cells (GC and ASC; CD27^+^CD38^+^) either cultured in 2D without FRCs, or 2D with FRCs, in RPMI versus IMDM, with or without additional supplementation of B cell cytokines IL4 and IL21. **(C)** Total counts of CD8^+^ T cells, naive CD4^+^ T cells (CD45RA^+^), Th1 T cells (CD4^+^CD45RA^-^ CXCR5^-^CXCR3^+^CCR6^-^), Th2 T cells (CD4^+^CD45RA^-^CXCR5^-^CXCR3^-^CCR6^-^), Th17 T cells (CD4^+^CD45RA^-^CXCR5^-^CXCR3^-^CCR6^+^), Th1/17 T cells (CD4^+^CD45RA^-^CXCR5^-^CXCR3^+^CCR6^+^) and Tfh T cells (CD4^+^CD45RA^-^CXCR5^+^) either cultured in 2D without FRCs, or 2D with FRCs, in RPMI versus IMDM, with or without additional supplementation of B cell cytokines IL4 and IL21. **(D)** Percentage (%) of viable lymphocytes when cultured in 2D without FRCs, 2D with FRCs, 3D without FRCs, or 3D with FRCs, comparing the effect of additional 1μg/mL recombinant human B cell-activating factor (BAFF) supplementation in IMDM medium without IL4 and IL21 supplementation. Data showing the mean ± SD (n=4 tonsil donors).

**Figure S3. Impact of FRCs and 3D culture on B and T cell survival, differentiation, and immunoglobulin production in response to viral antigens.** Survival of B and T cells when cultured in 2D without FRCs, 2D with autologous FRCs, or 3D with autologous FRCs, either left unstimulated or stimulated with SARS-CoV-2 WT spike protein (with and without R848), SARS-CoV-2 WT-I53-50 nanoparticles (NP), influenza H1N1 HA protein (with and without R848), influenza vaccine 2022/2023 (Influvac Tetra 2022/2023) or whole inactivated recombinant A/Netherlands/602/2009 influenza virus. Total viable B and T cell numbers are analyzed, as well as differentiation of B and T cells and total immunoglobulin production. **(A)** Viable CD19^+^ B cell counts on day 14 of each tested culture condition. **(B)** Viable CD4^+^ T cell counts on day 14 of each tested culture condition. **(C)** Total counts of double-negative cells (DN; CD27^-^CD38^-^IgD^-^), naive B cells (CD27^-^CD38^-^IgD^+^), memory B cells (CD27^+^CD38^-^), pre-GC (CD27^-^CD38^+^), and GC and antibody-secreting cells (GC and ASC; CD27^+^CD38^+^) either cultured in 2D without FRCs (left), 2D with FRCs (middle) or 3D with FRCs (right) cultured unstimulated, or for the varying tested antigen conditions at day 14. **(D)** Total immunoglobulin (Ig) production (IgM, IgG, and IgA) in the supernatant of tonsil cultures on day 14. Cultures were in 2D without FRCs (left), 2D with FRCs (middle), or 3D with FRCs (right), either unstimulated or stimulated with the various antigen conditions. Ig concentration is in ng/mL. **(E)** Total counts of naive CD4^+^ T cells (CD45RA^+^), Th1 T cells (CD4^+^CD45RA^-^CXCR5^-^CXCR3^+^CCR6^-^), Th2 T cells (CD4^+^CD45RA^-^CXCR5^-^CXCR3^-^CCR6^-^), Th17 T cells (CD4^+^CD45RA^-^CXCR5^-^CXCR3^-^CCR6^+^), Th1/17 T cells (CD4^+^CD45RA^-^CXCR5^-^ CXCR3^+^CCR6^+^) and Tfh T cells (CD4^+^CD45RA^-^CXCR5^+^) either cultured in 2D without FRCs (left), 2D with FRCs (middle), or 3D with FRCs (right) cultured unstimulated, or for the varying tested antigen conditions at day 14. (A and B) Data showing the mean (n=5 tonsil donors), (C-E) Data showing the mean ± SD (n=5 tonsil donors). *=p<0.05, **=p<0.01, ***=p<0.001, ****=p<0.0001, statistical significance is displayed in the color of the corresponding analyzed experimental group.

**Figure S4. Influence of FRCs and culture dimensionality on B cell survival.** Survival of B cells when cultured in 2D without FRCs, 2D with autologous FRCs, 2D with allogeneic FRCs, 3D with autologous FRCs or 3D with allogeneic FRCs, either left unstimulated or stimulated with SARS-CoV-2 WT spike (with and without R848), SARS-CoV-2 WT-I53-50 nanoparticles (NP), influenza H1N1 HA (with and without R848), influenza vaccine 2022/2023 (Influvac Tetra 2022/2023) or whole inactivated recombinant A/Netherlands/602/2009 influenza virus. Total viable B cell counts are shown for each individual donor tested, in all studied culture conditions, at day 14.

**Figure S5. Analysis of influenza- and SARS-CoV-2-specific B cell responses and bystander activation in tonsil co-cultures.** Characterization of both influenza H1N1 hemagglutinin (HA)-specific B cell responses and SARS-CoV-2 wild type (WT) spike-specific B cell responses after culture. Percentages of protein-specific cells are analyzed using flow cytometry. Protein-specific antibody responses were quantified using a Luminex assay, simultaneously quantifying protein unspecific bystander activation. **(A)** Percentage (%) of influenza H1N1 HA-specific B cells out of total living CD19^+^ B cells on day 14 for tonsil donors 1, 4, and 9. The percentage of influenza HA-specific B cells were quantified using flow cytometry in unstimulated cultures, or cultures stimulated with either influenza H1N1 HA-protein (with and without R848), influenza vaccine 2022/2023 (Influvac Tetra 2022/2023) or whole inactivated recombinant A/Netherlands/602/2009 influenza virus. All cultures were performed either in 2D without FRCs, 2D with autologous FRCs, 2D with allogeneic FRCs, 3D with autologous FRCs, or 3D with allogeneic FRCs. **(B)** Percentage (%) of SARS-CoV-2 WT spike-specific B cells out of total living CD19^+^ B cells on day 14 for tonsil donors 1, 4, and 9. The percentage of spike-specific B cells was quantified using flow cytometry in an unstimulated culture, or cultures stimulated with either SARS-CoV-2 WT spike (with and without R848) or SARS-CoV-2 WT-I53-50 NPs. All cultures were performed either in 2D without FRCs, 2D with autologous FRCs, 2D with allogeneic FRCs, 3D with autologous FRCs, or 3D with allogeneic FRCs. **(C)** Influenza H1N1 HA-specific IgM production (mean fluorescence intensity (MFI minus blank)) in the supernatants of the 2D and 3D (co-)cultures for tonsil donor 1, 4, and 9, for all culture conditions and antigen stimulations tested. **(D)** SARS-CoV-2 WT receptor-binding domain (RBD)-specific IgM production (mean fluorescence intensity (MFI minus blank)) in the supernatants of the 2D and 3D (co-)cultures for tonsil donor 1, 4, and 9, for all culture conditions and antigen stimulations tested. **(E)** Influenza H1N1 HA-specific IgG production (mean fluorescence intensity (MFI minus blank)) in the supernatants of the 2D and 3D (co-)cultures for tonsil donor 1, 4, and 9, for all culture conditions and antigen stimulations tested. **(F)** SARS-CoV-2 WT receptor-binding domain (RBD)-specific IgG production (mean fluorescence intensity (MFI minus blank)) in the supernatants of the 2D and 3D (co-)cultures for tonsil donor 1, 4, and 9, for all culture conditions and antigen stimulations tested. **(G)** Tetanus toxoid-specific IgG production (mean fluorescence intensity (MFI minus blank)) in the supernatants of the 2D and 3D (co-)cultures for tonsil donor 6, 8, and 9, for all culture conditions and antigen stimulations tested. **(H)** Influenza H1N1 HA-specific IgG production (mean fluorescence intensity (MFI minus blank)) in the supernatants of the 2D and 3D (co-)cultures for tonsil donor 6, 8, and 9, for all culture conditions and antigen stimulations tested. **(I)** SARS-CoV-2 WT receptor-binding domain (RBD)-specific IgG production (mean fluorescence intensity (MFI minus blank)) in the supernatants of the 2D and 3D (co-)cultures for tonsil donor 6, 8, and 9, for all culture conditions and antigen stimulations tested. **(J)** SARS-CoV-2 WT spike-specific IgG production (mean fluorescence intensity (MFI minus blank)) in the supernatants of the 2D and 3D (co-)cultures for tonsil donor 6, 8, and 9, for all culture conditions and antigen stimulations tested. Data showing the mean of technical duplicates (n=2) from the donors as indicated (A-F).

**Figure S6. Gating strategy for germinal center B cells and ASCs and their surface CXCR4/CXCR5 chemokine expression. (A)** Representative flow cytometry plot, showing the gating strategy used to identify germinal center (GC) B cells (CD20^+^CD38^+^) and antibody-secreting cells (ASCs, CD20^-^CD38^++^) out of total living CD19^+^ B cells. **(B)** Percentage (%) of mature B cell subsets; germinal center (GC) B cells (CD20^+^CD38^+^) and antibody-secreting cells (ASCs, CD20^-^CD38^++^), out of total living CD19^+^ B cells on day 14, comparing 2D without FRCs, 2D with autologous FRCs and 3D with autologous FRCs for both the low and high responsive tonsil donor. **(C)** Surface CXCR4 and CXCR5 chemokine expression of B cells cultured in 2D without FRCs, 2D with autologous FRCs, or 3D with autologous FRCs, either left unstimulated or stimulated with SARS-CoV-2 WT spike protein (with and without R848), SARS-CoV-2 WT-I53-50 NP, influenza H1N1 HA protein (with and without R848), influenza vaccine 2022/2023 (Influvac Tetra 2022/2023) or whole inactivated recombinant A/Netherlands/602/2009 influenza virus. Top row: total living CD19^+^ B cells, middle row: total living CD19^+^CD20^+^CD38^+^ B cells (GC B cells), bottom row: total living CD19^+^CD20^-^CD38^++^ B cells (ASCs). **(D)** Flow cytometry plots showing the expression of CD20 and BCL6 in relation to CD38 and CD27, or the expression of BCL6 in relation to CD38 and CD20, gated on viable CD19^+^ B cells. Data in (B) and (D) showing the mean ± SD (n=5 tonsil donors). *=p<0.05, **=p<0.01, ***=p<0.001, ****=p<0.0001, statistical significance is displayed in the color of the corresponding analyzed experimental group.

**Figure S7. Extended comparison of tonsil cell responses in 2D and 3D cultures with or without autologous or allogeneic FRCs.** Shown are 2D cultures with autologous or allogeneic FRCs and 3D cultures with autologous or allogeneic FRCs. **(A)** Expression of activation markers OX40 and CD69 on viable CD4⁺CD45RA⁻ T cells at day 7 (left) and day 14 (right) in unstimulated conditions. **(B)** Total counts of CD8⁺ T cells, naïve CD4⁺ T cells (CD45RA⁺), Th1 (CD4⁺CD45RA⁻CXCR5⁻CXCR3⁺CCR6⁻), Th2 (CD4⁺CD45RA⁻CXCR5⁻CXCR3⁻CCR6⁻), Th17 (CD4⁺CD45RA⁻CXCR5⁻CXCR3⁻CCR6⁺), Th1/17 (CD4⁺CD45RA⁻CXCR5⁻CXCR3⁺CCR6⁺), and Tfh (CD4⁺CD45RA⁻CXCR5⁺) T cells at day 14. **(C)** Total counts of CD19^+^ B cells, including double-negative (DN; CD27⁻CD38⁻IgD⁻), naïve (CD27⁻CD38⁻IgD⁺), memory (CD27⁺CD38⁻), pre-GC (CD27⁻CD38⁺), and GC/antibody-secreting cells (GC and ASC; CD27⁺CD38⁺) after 14 days of culture, either unstimulated or stimulated with the indicated antigens. **(D)** Percentage of SARS-CoV-2 WT spike-specific B cells (left) and influenza H1N1 HA-specific B cells (right) within total living CD19⁺ B cells on day 14. **(E)** Total immunoglobulin levels (IgM, IgG, and IgA) in culture supernatants on day 14, quantified in ng/mL by ELISA. **(F)** SARS-CoV-2 WT RBD-specific IgG (left) and influenza H1N1 HA-specific IgG (right) in supernatants on day 14, measured by Luminex; MFI values are background-subtracted. Data are shown as mean ± SD (n = 5 tonsil donors). *p < 0.05, **p < 0.01, ***p < 0.001, ****p < 0.0001; statistical significance is indicated in the color of the corresponding experimental group.

