## Supplementary figures and images for "Enhanced human antigen-specific B cell responses using *in vitro* 3D tonsil cultures containing stromal cells"

### Supplemental Figures

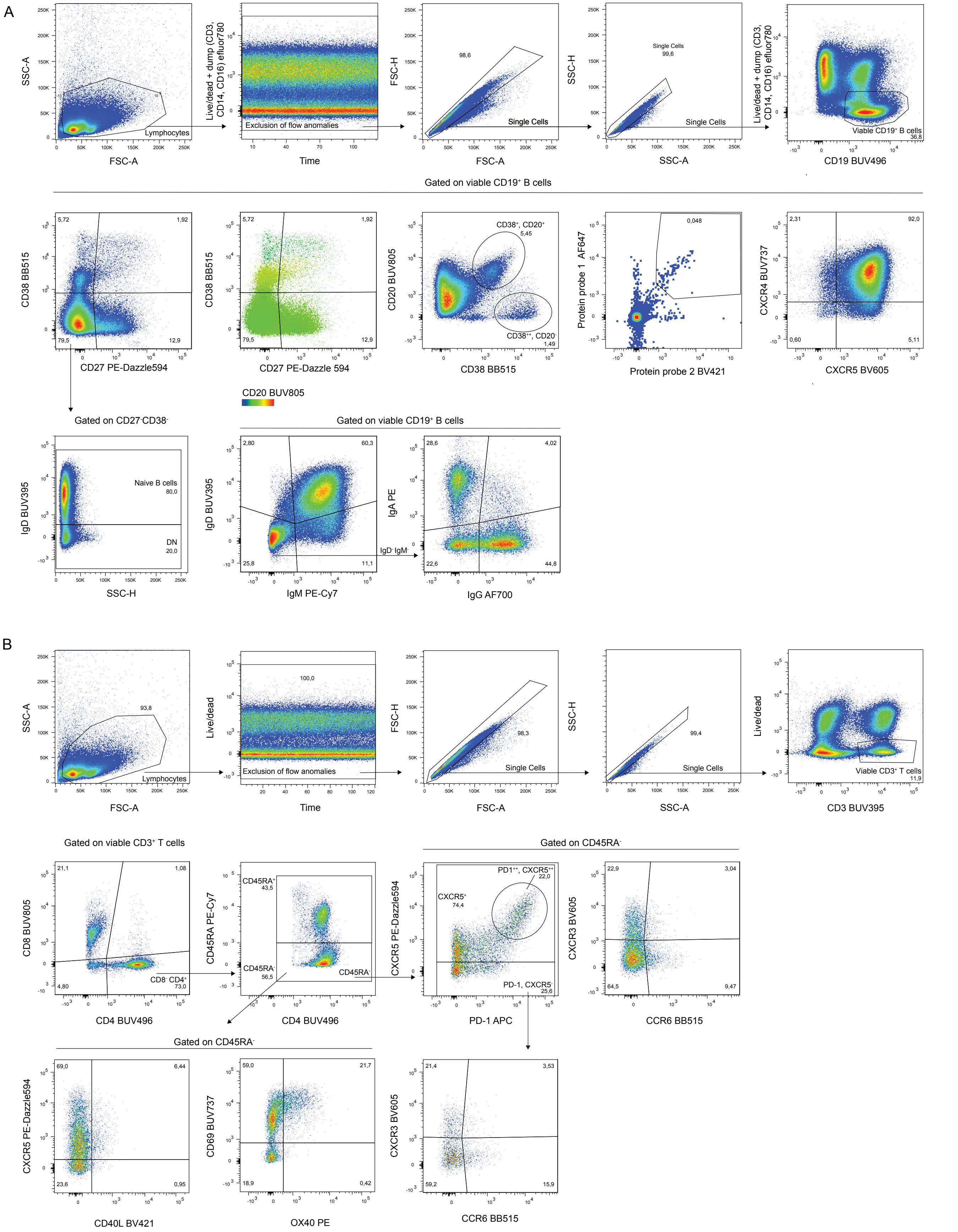

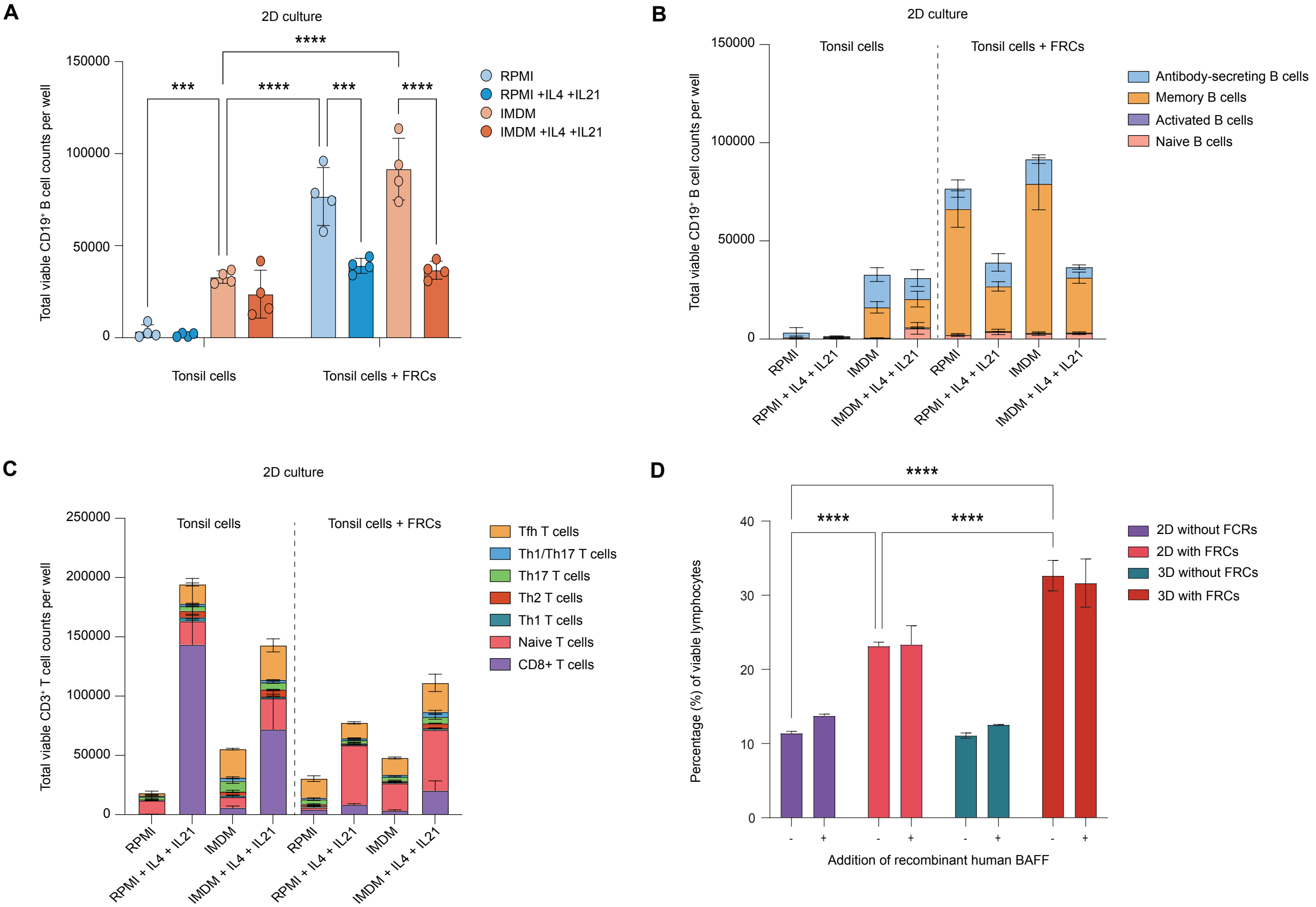

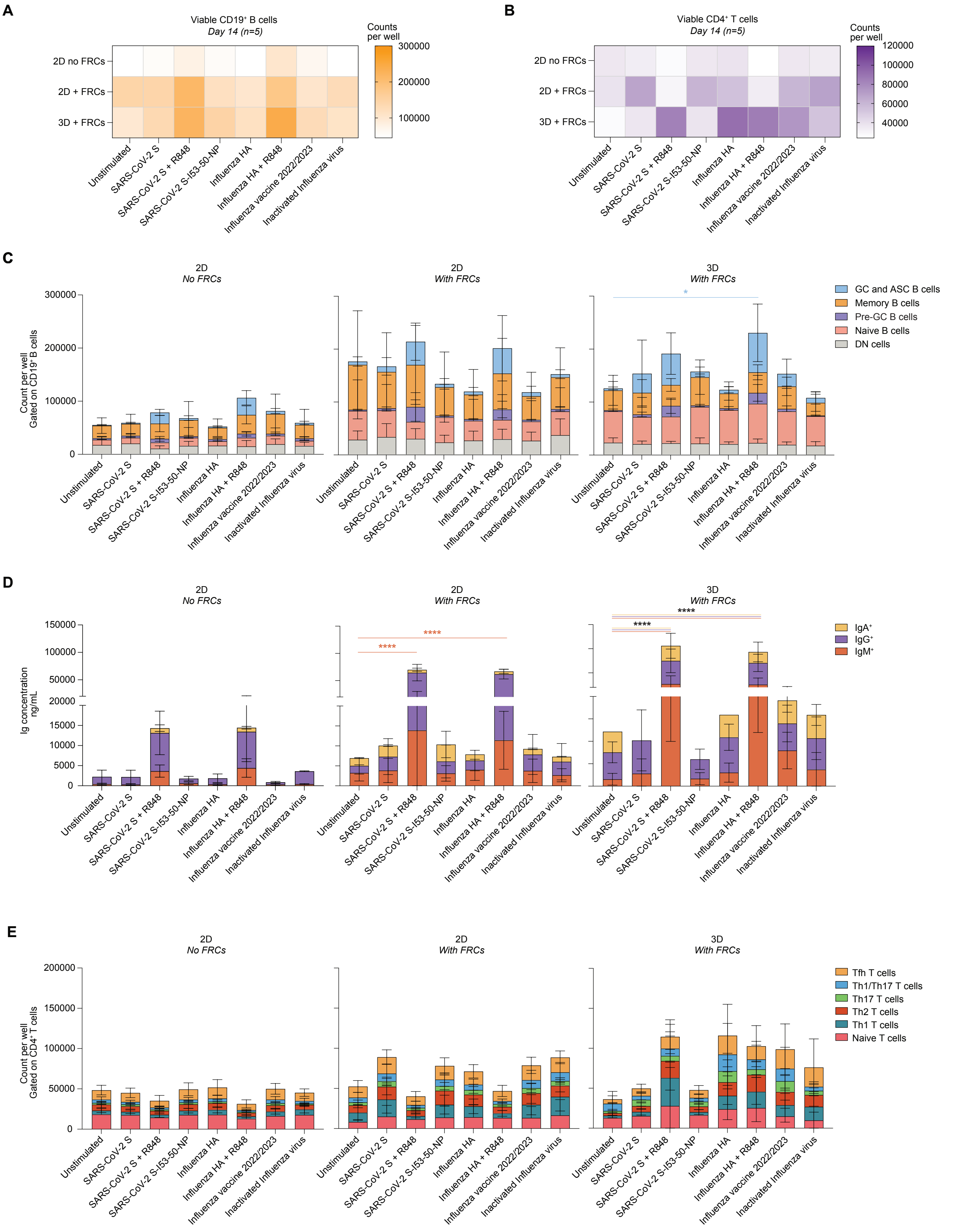

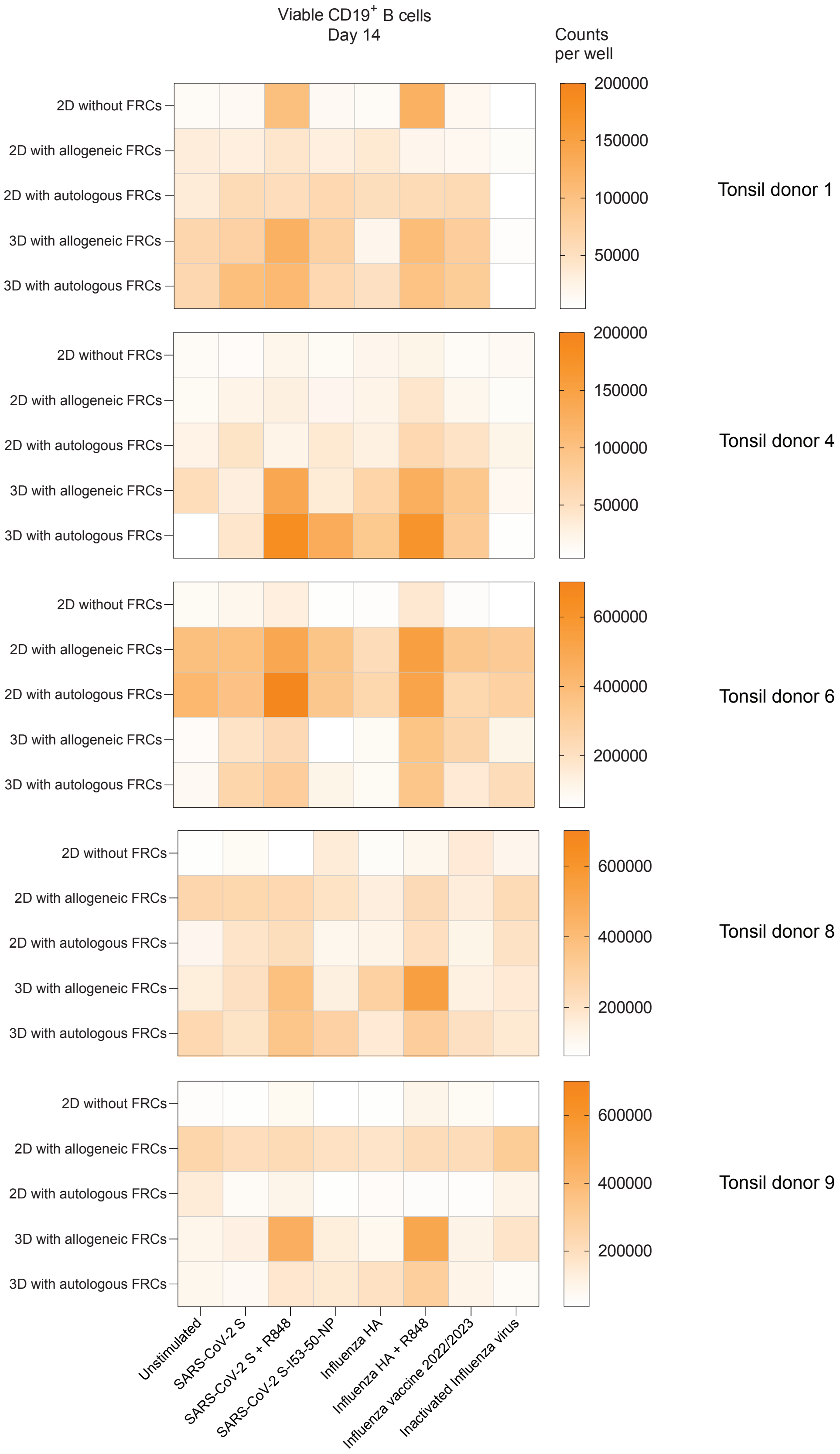

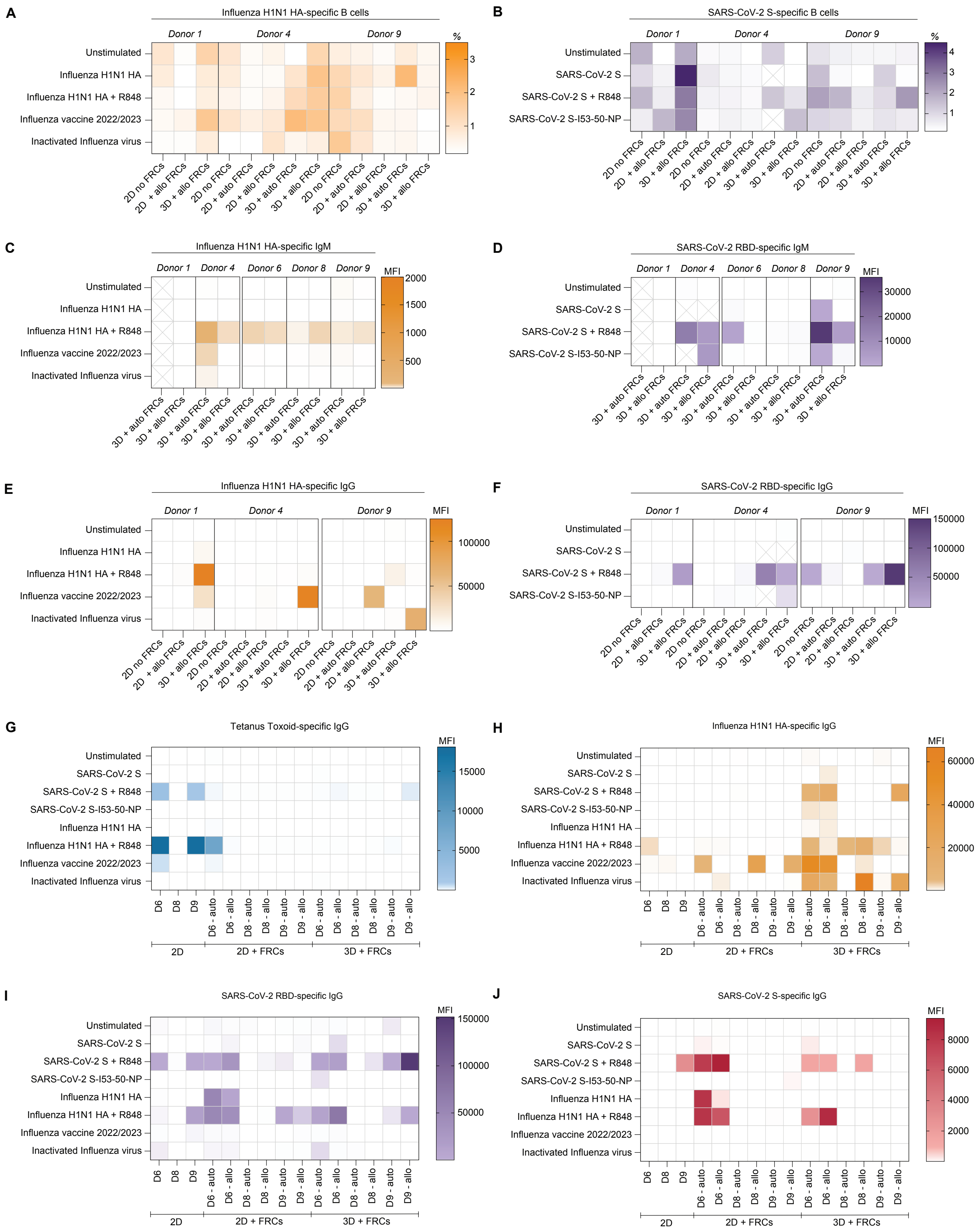

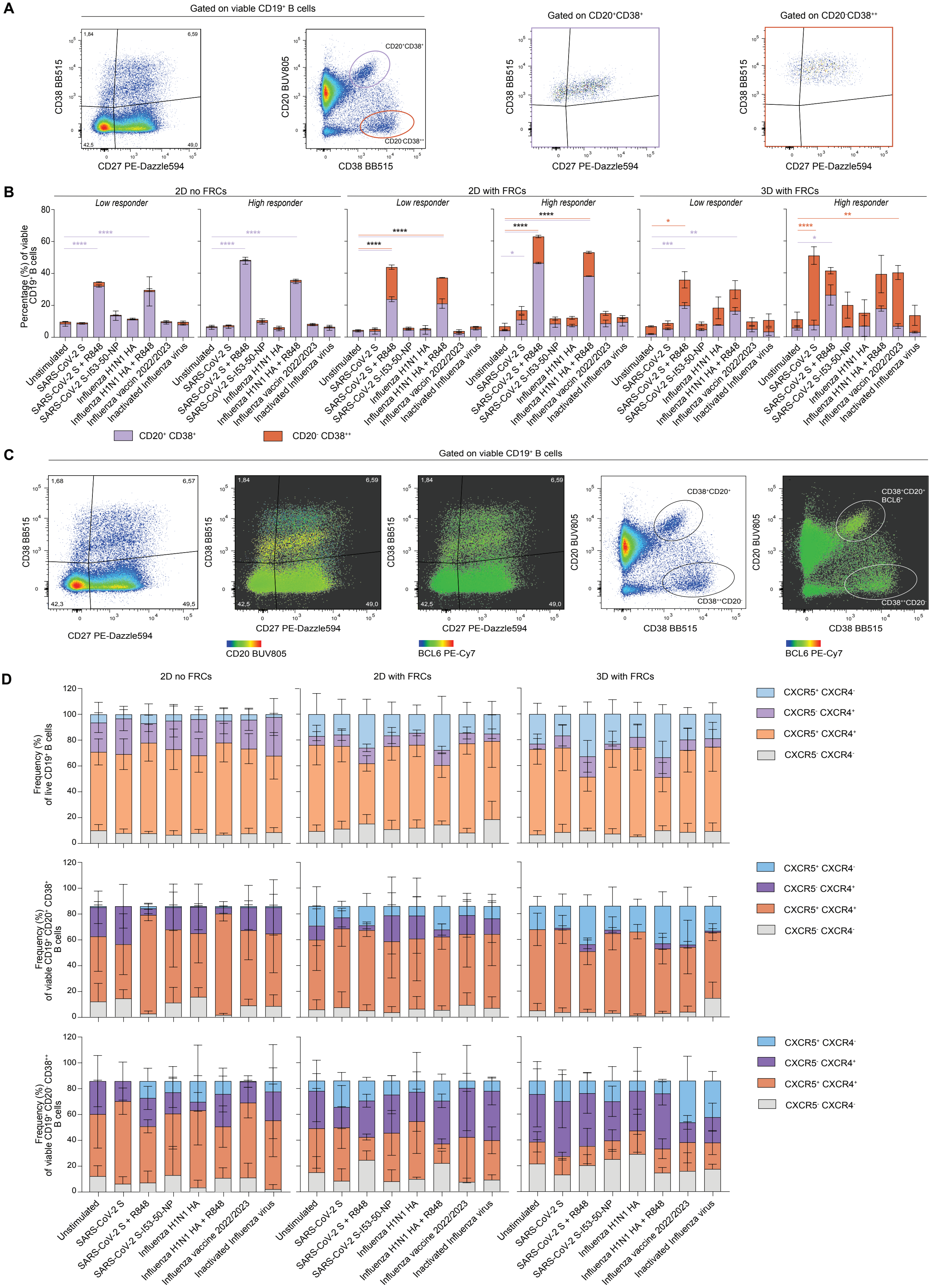

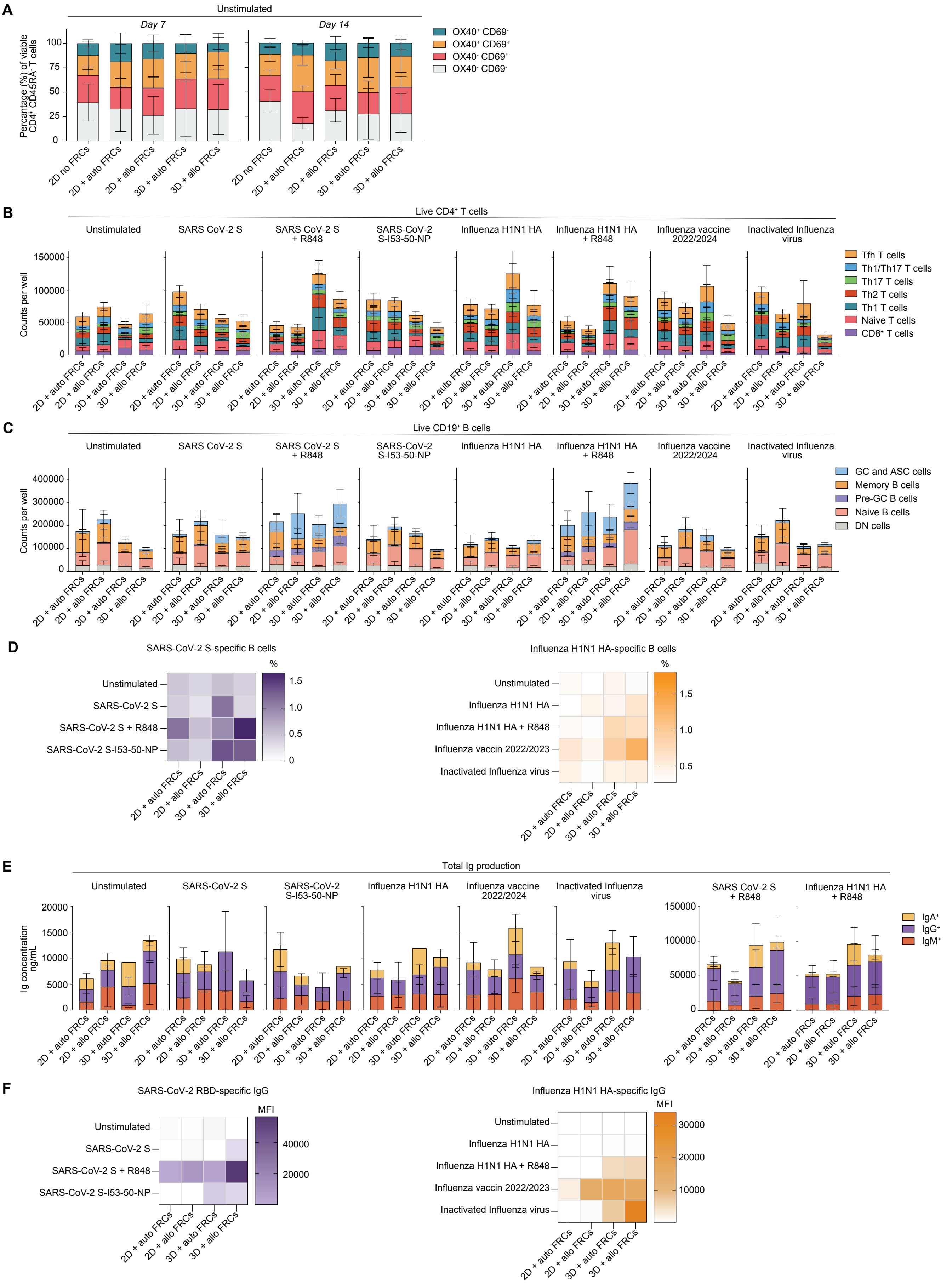
